# Genome-wide *in vivo* CRISPR screens identify GATOR1 as a potent tumor suppressor in MYC-driven lymphomagenesis

**DOI:** 10.1101/2022.02.16.480657

**Authors:** Margaret A. Potts, Shinsuke Mizutani, Yexuan Deng, Srimayee Vaidyanathan, Keziah E. Ting, Göknur Giner, Shruti Sridhar, Yang Liao, Sarah Diepstraten, Andrew J. Kueh, Martin Pal, Geraldine Healey, Lin Tai, Zilu Wang, Christina König, Deeksha Kaloni, Lauren Whelan, Michael Milevskiy, Hannah Coughlan, Giovanna Pomilio, Andrew H. Wei, Jane E. Visvader, Anthony T. Papenfuss, Stephen Wilcox, Anand D. Jeyasekharan, Wei Shi, Emily J. Lelliott, Gemma L. Kelly, Kristin K. Brown, Andreas Strasser, Marco J. Herold

## Abstract

Identifying tumor suppressor genes is predicted to inform on the development of novel strategies for cancer therapy. To identify new lymphoma driving processes that cooperate with oncogenic MYC (which is abnormally highly expressed in ∼70% of human cancers) we have used a genome-wide CRISPR knockout screen in *Eµ-Myc;Cas9* transgenic hematopoietic stem and progenitor cells *in vivo*. We discovered that loss of any of the GATOR1 complex components - NPRL3, DEPDC5, NPRL2 - significantly accelerated c-MYC-driven lymphoma development in mice. Low expression of the GATOR1 complex genes correlated with poor survival outcomes for human patients with high MYC-expressing cancers. Murine lymphomas lacking GATOR1 were highly sensitive to mTOR inhibitors as a single agent therapy, both *in vitro* and *in vivo*. These findings identify inhibition of mTORC1 as a potent tumor suppressive mechanism in c-MYC-driven lymphomagenesis and suggest a new avenue for therapeutic intervention in GATOR1-deficient lymphomas through mTOR inhibition.

**Key Points:**

- *In vivo* CRISPR/Cas9 whole genome knockout screens identified the GATOR1 complex as a potent suppressor of c-MYC-driven lymphomagenesis.
- GATOR1 deficiency in MYC-driven lymphomas confers sensitivity to mTOR inhibition, suggesting a novel therapeutic approach.

## Introduction

Abnormally high expression of the oncogene *c-MYC* is observed in ∼70% of human cancers, highlighting it as an attractive therapeutic target ^1^. However, to date no MYC directed therapeutics have been successfully developed. Many cellular processes are regulated by MYC, including cell cycle control and proliferation, differentiation, metabolism and cell death ^2^. Understanding which processes are critical for MYC-driven oncogenesis might reveal downstream druggable cancer dependencies.

To identify genes that suppress MYC-driven cancer, we conducted an unbiased genome-wide CRISPR/Cas9 knockout screen *in vivo*. We employed the *Eµ-Myc* transgenic mouse model that serves as a valuable pre-clinical model of Burkitt Lymphoma (BL) and other MYC-driven malignancies. These mice were engineered such that human c-MYC is constitutively expressed at high levels throughout B cell development under the control of the murine *Eµ*-enhancer. Consequently, this induces a partial differentiation block and increased expansion of early B cell progenitors that amass as a pool of pre-leukemic cells ^3,4^. These pre-leukemic cells are prone to undergoing apoptosis, and acquisition of additional cooperating mutations enables malignant transformation into monoclonal surface Ig- pre-B lymphoma or surface Ig+ B-cell lymphoma that disseminates to all hematopoietic organs with mice succumbing to disease with a median survival of ∼110 days ^5^.

While genome-wide CRISPR/Cas9 screening approaches are feasible *in vitro* using immortalized or easily transducible cell lines, they do not accurately model the complexities of cancer initiation, metastasis, nor response to therapies or interactions within the tumor microenvironment ^6,7^. Previously, only few *in vivo* CRISPR/Cas9 screens have employed non-transformed cells to interrogate cancer initiation ^8–10^. Moreover, most such *in vivo* screens exploring tumorigenesis were performed either with small focused sgRNA ^11,12^ or siRNA ^13^ libraries, thereby limiting the hit identification to known cellular signaling pathways. Other screens employed genome-wide targeting, albeit by using transplantation of established cancer cell lines, which possess complex (and possibly confounding) genetic aberrations, rather than primary cells.

Here, we conducted an unbiased genome-wide CRISPR/Cas9 knockout screen in primary cells *in vivo* to identify tumor suppressor genes whose loss cooperates with oncogenic MYC in driving lymphomagenesis. Primary *Eµ-Myc;Cas9* double transgenic hematopoietic stem and progenitor cells (HSPCs) that were transduced with a genome-wide sgRNA library were transplanted into lethally irradiated mice. The transplanted HSPCs can reconstitute all blood compartments while also being predisposed to developing lymphoma, whereby CRISPR/Cas9 deletion of a tumour suppressor gene would induce malignant transformation. Since lymphomas in the *Eµ-Myc* transgenic model are monoclonal ^5^, our screen will identify the most powerful tumor suppressor pathways as each manipulated *Eµ-Myc* transgenic HSPC deleted for one gene and transplanted into a recipient mouse will have to compete with thousands of other genetically manipulated HSPCs, resulting in the selection of the strongest hit in each recipient mouse.

Aberrations in regulators of the cell death pathway are well known to cooperate with oncogenic c-MYC in lymphomagenesis ^14–21^. This was evident by the observation that deletion of the tumour suppressor gene p53 was the most frequent hit in our screen. However, several top hits were genes involved in the negative regulation of the mTORC1 pathway, which coordinates cellular metabolism to support cell growth, survival and proliferation ^22^. Two components of the mTORC1 inhibitory complex GATOR1 were top hits and they were not previously described as suppressors of MYC driven lymphomagenesis. We revealed the importance of this negative regulator of cell metabolism for tumor suppression, including in human cancers through analysis of human cancer genome databases. Of note, we show that lymphomas driven by loss of GATOR1 are highly sensitive to single agent mTOR inhibitor treatment, suggesting a therapeutic strategy for cancers with oncogenic MYC that are also GATOR1-deficient.

## Results

### Genome-wide *in vivo* CRISPR/Cas9 gene knockout screen identifies mTOR inhibitory pathway genes as strong suppressors of MYC-driven lymphoma

Previously our laboratory conducted an RNA interference (RNAi) screen where the siRNA library was focused on p53 regulated genes ^13^. To identify novel suppressors of c-MYC driven lymphomagenesis beyond the p53 pathway we performed an unbiased CRISPR/Cas9 gene knockout screen *in vivo*. Specifically, fetal liver cells, an abundant source of hematopoietic stem/progenitor cells (HSPCs), from C57BL/6-Ly5.2 *Eµ-Myc;Cas9* double transgenic day (E)13.5 mouse embryos were transduced *ex vivo* with a pooled whole-genome sgRNA library. The library contains 87,987 sgRNAs targeting 19,150 mouse protein coding genes (4-5 sgRNAs per gene) ^23^ and was transduced at an efficiency of 20-30%, minimizing the likelihood of co-transducing multiple sgRNAs into a single cell. At 24 h post-transduction, the HSPCs were transplanted intravenously into lethally irradiated congenic C57BL/6-Ly5.1 recipient mice (Fig. 1A). As a positive control, *Eµ-Myc;Cas9* HSPCs were transduced with a sgRNA targeting mouse *p53* (*sgp53)*, as loss of *p53* is known to substantially accelerate c-MYC-driven lymphomagenesis ^24^. As a negative control, *Eµ-Myc;Cas9* HSPCs were transduced with a sgRNA targeting the human *BIM* gene that has no predicted activity in the mouse genome (*sgControl*). Recipient mice transplanted with *sgControl* HSPCs developed lymphoma spontaneously with a median latency of ∼140 days post-transplantation, while those transplanted with *sgp53* HSPCs demonstrated significantly accelerated lymphoma development (median latency of ∼25 days) (Fig. 1B), consistent with a previous report ^25^. Notably, several mice transplanted with sgRNA library-transduced HSPCs developed lymphoma considerably earlier (median latency ∼74 days) than the mice transplanted with *sgControl* HSPCs (Fig. 1B). This indicated that these accelerated lymphomas contained sgRNAs that led to the deletion of genes involved in the suppression of c-MYC-driven lymphomagenesis. All transplanted mice, regardless of the sgRNA or disease onset, had sIg^-^ pre-B or sIg^+^ B cell lymphoma (Supplemental Fig. S1A-B), as expected in this model ^5^.

**Figure 1.**
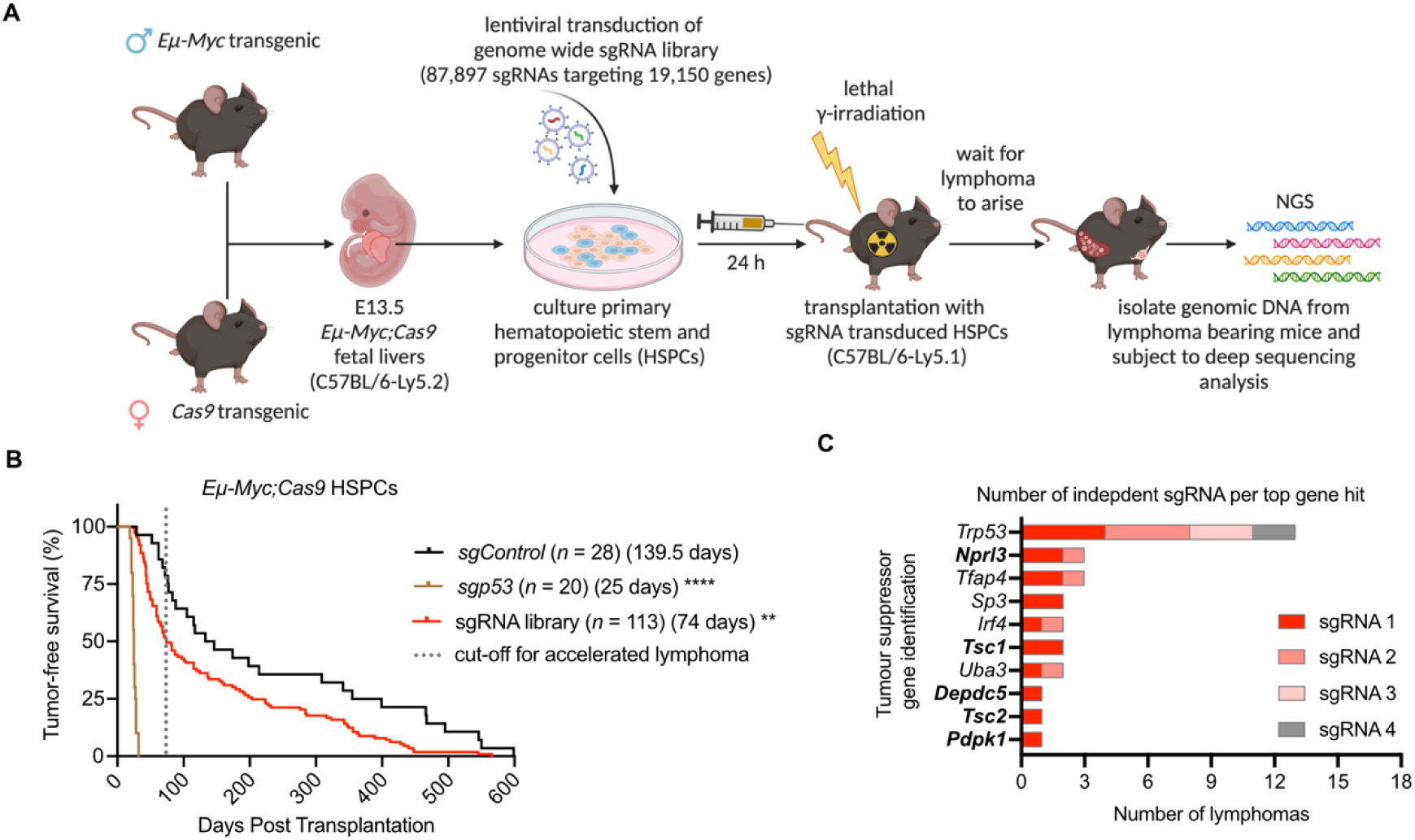
*In vivo* genome-wide CRISPR/Cas9 gene knockout screen identifies novel candidate tumor suppressors. **A**, Schematic of the experimental strategy for performing *in vivo* genome-wide sgRNA screens to identify novel tumor suppressors. Fetal liver cells, a rich source of hematopoietic stem/progenitor cells (HSPCs), from E13.5 *Eµ-Myc*;*Cas9* mice (C57BL/6-Ly5.2 background) were transduced with lentiviruses containing sgRNAs targeting *p53* (*sgp53;* positive control), a negative control sgRNA targeting human *BIM* (*sgControl)* or a whole-genome sgRNA library ^23^. The transduced HSPCs were injected intravenously (i.v.) into lethally irradiated (2x5.5 Gy, 3 h apart) congenic recipient C57BL/6-Ly5.1 mice. Lymphoma bearing mice displayed enlarged spleen, lymph nodes and/or thymus. These tissues were harvested and cryopreserved. Genomic DNA was isolated from the spleen, comprising mostly of lymphoma cells but also containing non-transformed hematopoietic cells; sgRNAs were identified by NGS. **B,** Tumor-free survival of mice transplanted with *Eµ-Myc;Cas9* HSPCs that had been lentivirally transduced with the positive control sgRNA (*sgp53)*, the negative control sgRNA (*sgControl)* or a whole-genome sgRNA library. The dotted line represents cut-off for lymphomas arbitrarily deemed to be accelerated for further analysis. n represents total number of transplanted mice per sgRNA from 6 reconstitution cohorts. Median survival is indicated in brackets. Log-rank (Mantel-Cox) statistical test for survival curve comparison, \*\**P*=0.0002, \*\*\*\**P < 0*.0001. **C,** Top 10 tumor suppressor genes identified, determined by frequency of their sgRNA detected in independent lymphomas from mice with accelerated lymphoma onset and being highly enriched (>50% of reads within a given lymphoma) by sequencing. Genes emboldened represent those in the mTORC1 inhibitory pathway of which five of the top 10 genes belong to this pathway.

To identify the sgRNAs that caused accelerated lymphomagenesis, we conducted next generation sequencing (NGS) on genomic DNA extracted from the spleens of mice transplanted with sgRNA library-transduced HSPCs that developed lymphoma prior to 75 days after reconstitution (arbitrary cut-off for accelerated lymphoma chosen; *n* = 60 from 113 transplanted mice) (Fig. 1B). Analysis of gDNA from whole spleen, which consists of both malignant lymphoma cells and non-malignant blood cells revealed one dominant sgRNA accounting for a large proportion of the sequencing counts, which we designated as the candidate tumor suppressor gene (Figure 1C, Supplemental Fig. S2A). Additionally, we observed low read counts of numerous sgRNAs collectively targeting more than 75% of all genes. These sgRNAs likely conferred no selection pressure and are present in the non-malignant hematopoietic cells within the spleens of the transplanted mouse (Supplemental Fig. S2B-F). This provides an important internal control to demonstrate high level sgRNA library representation in each reconstituted mouse. In fact, of the genes for which no sgRNA was represented (∼25%), 809 (∼4%) are considered essential genes according to DepMap and the TCGA, and therefore would not be expected to be represented in the reconstituted mice^26^. Collectively, these data indicate comprehensive gene coverage of the genome-wide sgRNA library that is usually significantly reduced following transplantation into mice ^27^.

Overall, 38 top hits were identified from the 60 lymphoma samples sequenced, with some sgRNAs targeting the same gene appearing in multiple samples, including several known tumor suppressors genes. The most prominent hit was *p53*, with four of the five sgRNAs targeting *p53* being the dominant sgRNA in 13 different lymphomas (Fig. 1C, Supplemental Fig. S2A). Notably, the screen also identified new candidate tumor suppressor genes, such as *Tfap4*, where two sgRNAs were dominant in three lymphomas. This gene was investigated in a separate study ^28^. Pathway analysis of the top hits using the DAVID Bioinformatics tool revealed that five of the top hits - *Nprl3*, *Depdc5*, *Tsc1*, *Tsc2* and *Pdpk1*- belong to the mTOR signaling pathway ^29,30^. Among these hits, *Nprl3* (with two sgRNAs dominant in three lymphomas) and *Depdc5* (with one sgRNA dominant in one lymphoma) (Fig. 1C, Supplemental Fig. S2A) were particularly interesting as they both encode essential components of the GATOR1 complex that negatively regulates mTORC1 in response to amino acid availability ^22^. While one study identified loss of function mutations in the GATOR1 complex components in some solid malignancies ^31^, it has not previously been reported to suppress MYC-driven hematological malignancies. In contrast Tuberous sclerosis complex 1 (*Tsc1*) and *Tsc2,* which also negatively regulate mTORC1, but in response to growth factor signaling, have previously been implicated in the suppression of MYC-driven lymphoma development ^22,32,33^. These findings indicate that our unbiased *in vivo* CRISPR gene knockout screening approach successfully selected for the strongest suppressors of MYC-driven lymphoma development, identifying both known as well as novel tumor suppressor genes. In particular, we found that the mTOR inhibitory pathway plays a previously underappreciated but critical role in MYC-driven lymphoma development as sgRNAs targeting two genes within this pathway outcompeted sgRNAs against all other genes.

### Loss of any GATOR1 complex component accelerates MYC-driven lymphoma development

Although only two out of three components of the GATOR1 complex were hits in our screen, all three – NPRL3, DEPDC5, and NRPL2 the third component of the GATOR1 complex (Fig. 2A) - are required for its function ^34^. Therefore, to validate NPRL3, DEPDC5 and NPRL2 as tumor suppressors, two independent sgRNAs targeting each of their genes were individually introduced into *Eµ-Myc;Cas9* HSPCs that were then transplanted into lethally irradiated recipient mice. Strikingly, all sgRNAs targeting *Nprl3*, *Depdc5* or *Nprl2* significantly accelerated c-MYC-driven lymphomagenesis to a similar extent as each other or even sgRNAs targeting the potent tumor suppressor p53 (sg*p53)* (Fig. 2B-D). As expected in the *Eµ-Myc* transgenic mouse model ^5^, all transplanted mice regardless of the sgRNA or lymphoma onset displayed enlarged hematopoietic tissues and elevated white blood cell counts at humane endpoint (Supplemental Fig. S3A-C). Efficient deletion of *Nprl3*, *Depdc5* or *Nprl2* by CRISPR/Cas9 was confirmed by targeted gene sequencing of gDNA isolated from cell lines derived from these lymphomas (Fig. 2E). These findings reveal that loss of any GATOR1 complex component is sufficient to abrogate its tumor suppressive function during MYC-driven lymphoma development.

**Figure 2.**
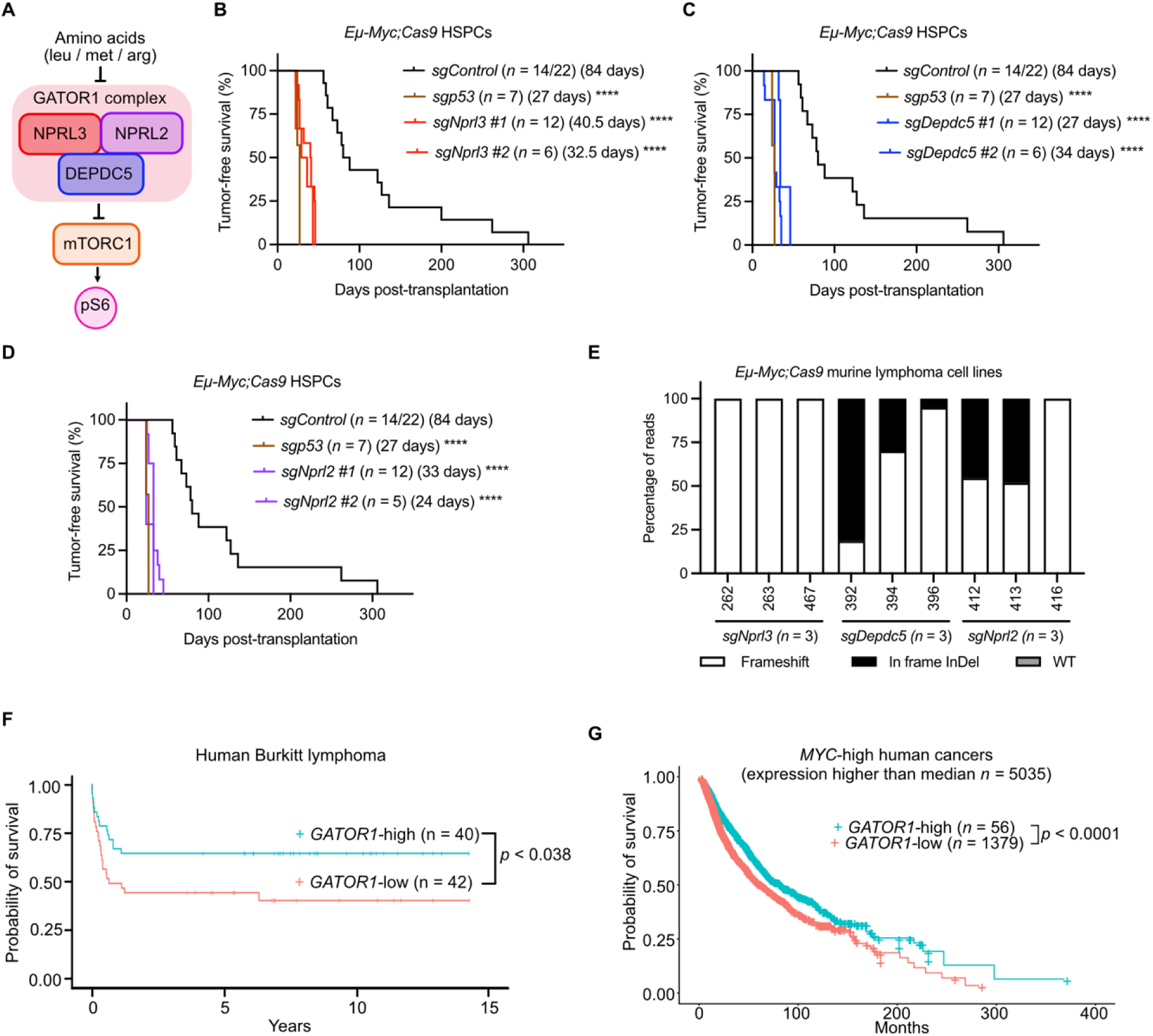
Validation of GATOR1 complex components as tumor suppressors of MYC driven lymphomagenesis in mice as well as in human MYC-driven cancers. **A**, Schematic of the GATOR1 complex, consisting of three proteins, NPRL3, DEPDC5 (both identified as hits in our genome-wide CRISPR screen), and NPRL2. The GATOR1 complex negatively regulates mTORC1 signaling in response to the availability of the amino acids leucine (leu), methionine (met) and arginine (arg). **B-D,** Tumor-free survival of mice transplanted with *Eµ-Myc;Cas9* HSPCs that had been transduced with either the sg*p53* (positive control), the negative control sgRNA (*sgControl*) or two independent sgRNAs each for targeting either *Nprl3* B, *Depdc5* C, or *Nprl2* D. *n* represents the number of transplanted mice per sgRNA across two transplanted mouse cohorts. Median survival is indicated in brackets. Log-rank (Mantel-Cox) test was used for comparison of mouse survival curves, \*\*\*\**P < 0*.0001. **E**, Proportions of frameshift, in frame InDels or WT sequence reads for the target gene of each sgRNA, analyzed by NGS of cell lines derived from lymphomas of recipient mice that had been transplanted with *sgNrpl3, sgDepdc5* or *sgNprl2* transduced *Eµ-Myc;Cas9* HSPCs. **F**, Probability of survival of human patients with Burkitt lymphoma, a cancer driven by *c-MYC* chromosomal translocations, stratified by *GATOR1* expression, where the *GATOR1*-low (*n* = 42) strata is defined as expression of either the *NPRL3*, *DEPDC5* or *NPRL2* mRNA in the lowest quartile. The others were grouped into the *GATOR1*-high strata (*n* = 40). Kaplan-Meier statistical test, \*\*\*\**P* < 0.038. **G**, Human TCGA pan-cancer analysis of *MYC*-high mRNA (expression higher than median, *n* = 5,035 patient samples) tumors, showing the difference in probability of survival between *GATOR1*-low group samples (defined as tumors with *NPRL2* or *DEPDC5* or *NPRL3* expression in the bottom quartile, median survival 47 months, *n* = 1,379 patient samples) versus *GATOR1-*high group samples (defined as tumors with *NPRL2* and *DEPDC5* and *NPRL3* expression in the top quartile, median survival 70 months, *n* = 56 patient samples). Kaplan-Meier statistical test, \*\*\*\**P* < 0.0001.

To examine if these findings are relevant to human cancer, we analyzed a Burkitt lymphoma patient cohort; this is a c-*MYC* gene chromosomal translocation driven B cell malignancy similar to the *Eµ-Myc* mouse lymphomas. Burkitt lymphoma samples were stratified by the levels of GATOR1 component gene expression. Those in the *GATOR1*-low expression strata (defined as *NPRL3*, *DEPDC*5 or *NPLR2* mRNA expression in the bottom 25%, *n* = 42 samples) correspond with significantly poorer probability of survival in patients compared to those in the *GATOR1*-high strata (n = 40 samples) (Fig. 2F). The rationale for this sample stratification is that the GATOR1 complex is non-functional when it lacks any one of the three components. Previous reports did not explore whether elevated MYC expression and loss of the GATOR1 inhibition of mTORC1 collaborate across other cancer types. Therefore, we extended our analysis to the pan-cancer data from The Cancer Genome Atlas (TCGA). Firstly, we examined all cancers with high *MYC* mRNA expression (defined as expression above the mean) that were then stratified by *GATOR1* complex mRNA expression. Consistent with the Burkitt lymphoma analysis, this revealed that *GATOR1-low* tumors, defined as those with either *NPRL3*, *DEPDC5* or *NPRL2* mRNA expression in the bottom quartile, were associated with significantly poorer overall patient survival compared to *GATOR1- high* tumors, defined as those with all of *NPRL3*, *DEPDC5* and *NPRL3* mRNA expression above the mean (Fig. 2G). These findings identify the GATOR1 complex as a suppressor of MYC-driven tumorigenesis.

### Loss of GATOR1 obviates the pressure for p53 mutation in c-MYC-driven lymphomagenesis

Given that deletion of GATOR1 components accelerated lymphomagenesis to a similar extent as p53 deletion and since MYC-lymphomas are prone to select for aberrations in the p53 pathway, we examined p53 mutational status and function in *sgControl* and GATOR1-deficient lymphomas. Consistent with previous reports that spontaneous mutations in p53 arise in ∼30% of *Eµ-Myc* lymphomas ^15,18^, we found that ∼30% of *sgControl* lymphomas had acquired p53 pathway defects, as evidenced by characteristic expression of a stabilized mutant p53 protein and/or downstream p19^ARF^ which is increased when p53 function is lost ^15,18,35^ (Fig. 3A, Supplemental Fig. S4A). In contrast, no such abnormalities in p53 were found in 24 GATOR1-deficient *Eµ-Myc* lymphomas (Fig. 3A, Supplemental Fig. S4A). To functionally validate these findings, we treated lymphoma cell lines in culture with nutlin-3a, which induces apoptotic death in a p53-dependent manner ^36,37^. Three out of 10 *sgControl* lymphomas were resistant to nutlin-3a treatment consistent with the predicted presence of mutations in p53, whereas all 27 GATOR1-deficient lymphomas were highly sensitive to nutlin-3a indicating the presence of functional wildtype p53 and its downstream pathways (*n* = 27; Fig. 3B, Supplemental Fig. S4B). NGS confirmed that the three nutlin-3a resistant *sgControl* lymphomas had indeed acquired pathologic *p53* mutations, whereas all 27 (100%) GATOR1-deficient lymphomas maintained wild-type *p53* status (Fig. 3C). Extending this analysis to an independent cohort of human diffuse large B cell lymphoma (DLBCL) patient samples, 11/73 (∼15%) harbored non-synonymous mutations in *DEPDC5* and 9/73 (∼12%) harbored non-synonymous mutations in *NPRL2*, mutational data on *NPRL3* were not available (Fig. 3D). All but one of these GATOR1 mutant human DLBCL samples carried wildtype p53, corroborating our findings in mice that mutations in *p53* and GATOR1 complex genes are mutually exclusive (Fig. 3D). Together, these data suggest that the GATOR1 complex is a strong suppressor of MYC-driven lymphomagenesis, obviating the pressure to spontaneously acquire mutations in the p53 pathway.

**Figure 3.**
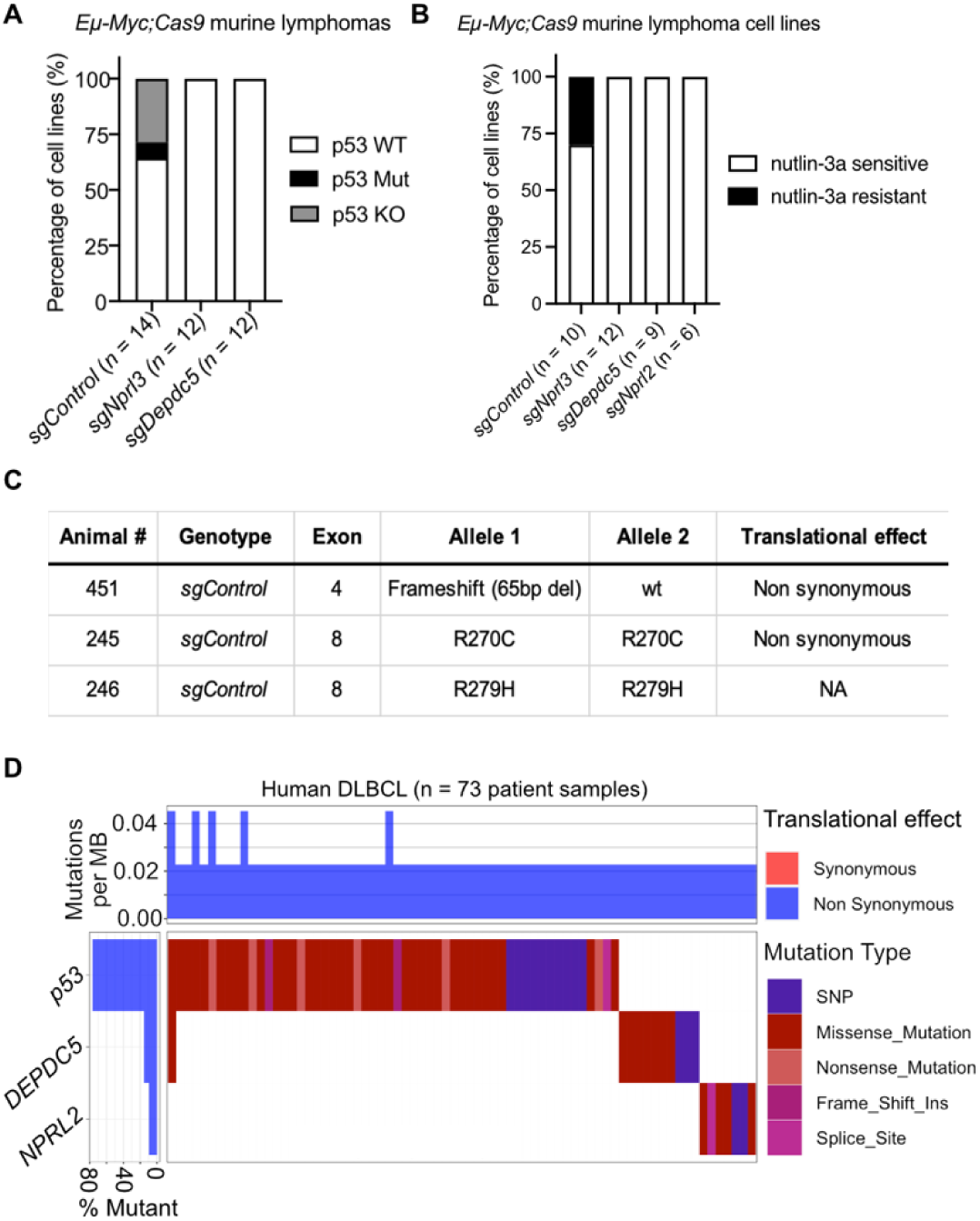
GATOR1 deficiency obviates the pressure to lose p53 function during c-MYC-driven lymphomagenesis. **A**, Summary graph of Western blot analyses showing proportions of *sgControl*, *sgDepdc5* or *sgNprl3 Eµ- Myc;Cas9* lymphomas that are p53 WT, p53 mutant (mut) or p53 knockout (KO). *n* = number of tumors for each genotype analyzed are indicated below the bars. **B**, Summary plot showing the proportions of nutlin-3a sensitive (p53 WT) or nutlin-3a resistant (p53 function defective) lymphoma cell lines per genotype. Each dot represents an independent experiment. **C**, Next generation sequencing of exons 4-11 of the *p53* genomic locus to identify mutations. 3/10 *sgControl*, 0/9 *sgNprl3*, 0/8 *sgDepdc5* and 0/7 *sgNprl2* lymphoma cell lines tested carried mutations in the *p53* gene. Translational effect determined from the equivalent human homologue mutation according to the TP53 database (accessed: https://tp53.cancer.gov/). **D**, Mutational analysis of *p53* and the GATOR1 component genes *DEPDC5* and *NPRL2* in human diffuse large B cell lymphoma (DLBCL) patient samples. Translational effect of mutation indicated, where synonymous refers to silent mutations and non-synonymous mutations encode an amino acid change. *n* = 73 patient samples, *P < 0.0001* using Fisher’s exact test.

### GATOR1 deficiency augments mTORC1 activity that can be diminished by direct inhibition of mTORC1

The GATOR1 complex is well established as an inhibitor of mTORC1 via specific amino acid sensing ^22^. We therefore hypothesized that lymphomas deficient in GATOR1 components would exhibit elevated phosphorylated S6, a surrogate marker of mTORC1 signaling, and fail to negatively regulate mTORC1 in response to amino acid availability. Indeed, at steady state, we observed substantially higher levels of phosphorylated S6 (phospho-S6; i.e. activated S6), indicative of increased mTORC1 activity in *Eµ-Myc* lymphoma cells lacking NPRL3 or DEPDC5 compared to control *Eµ-Myc* lymphoma cells (Fig. 4A-B). Even during culture in starvation medium (lacking all amino acids and serum) ^34^, phospho-S6 levels were higher in GATOR1-deficient *Eµ-Myc* lymphoma cells compared to control *Eµ-Myc*;Cas9 lymphoma cells (Fig. 4B). This emphasizes the loss of GATOR1 function, as the lack of amino acids should activate the GATOR1 complex thereby inhibiting mTORC1 (Fig. 4A). Although the magnitude of the elevation of phospho-S6 level above control in GATOR1-deficient *Eµ-Myc* lymphoma cells in starvation medium differed to that observed in steady state, this may be due to other amino acid sensors ^38^ dampen mTORC1 signaling and thus lower phospho-S6 levels as the starvation medium lacked all amino acids, not just the three sensed by GATOR1. In fact, starvation of the amino acids leucine (leu), methionine (met), and arginine (arg) that are uniquely sensed by the GATOR1 complex reduced phospho-S6 protein levels in control lymphomas (Fig. 4C; lane 3), however consistently, the GATOR1-deficient *Eµ-Myc* lymphomas failed to inhibit mTORC1 following amino acid starvation (Fig. 4C; lanes 7 and 11). Since GATOR1 deficient lymphomas displayed elevated mTORC1 activity, we considered whether mTORC1 inhibition would restore signaling to normal levels. Notably, treatment with rapamycin, a compound that directly inhibits mTORC1 (Fig. 4a), markedly reduced the levels of phospho-S6 in NPRL3 and DEPDC5 deficient lymphoma cells in both complete medium and starvation medium lacking all amino acids or only the amino acids sensed by the GATOR1 complex specifically (Fig. 4B-C). These findings demonstrate that the loss of GATOR1 function drives constitutive mTORC1 activation in *Eµ-Myc* lymphomas.

**Figure 4.**
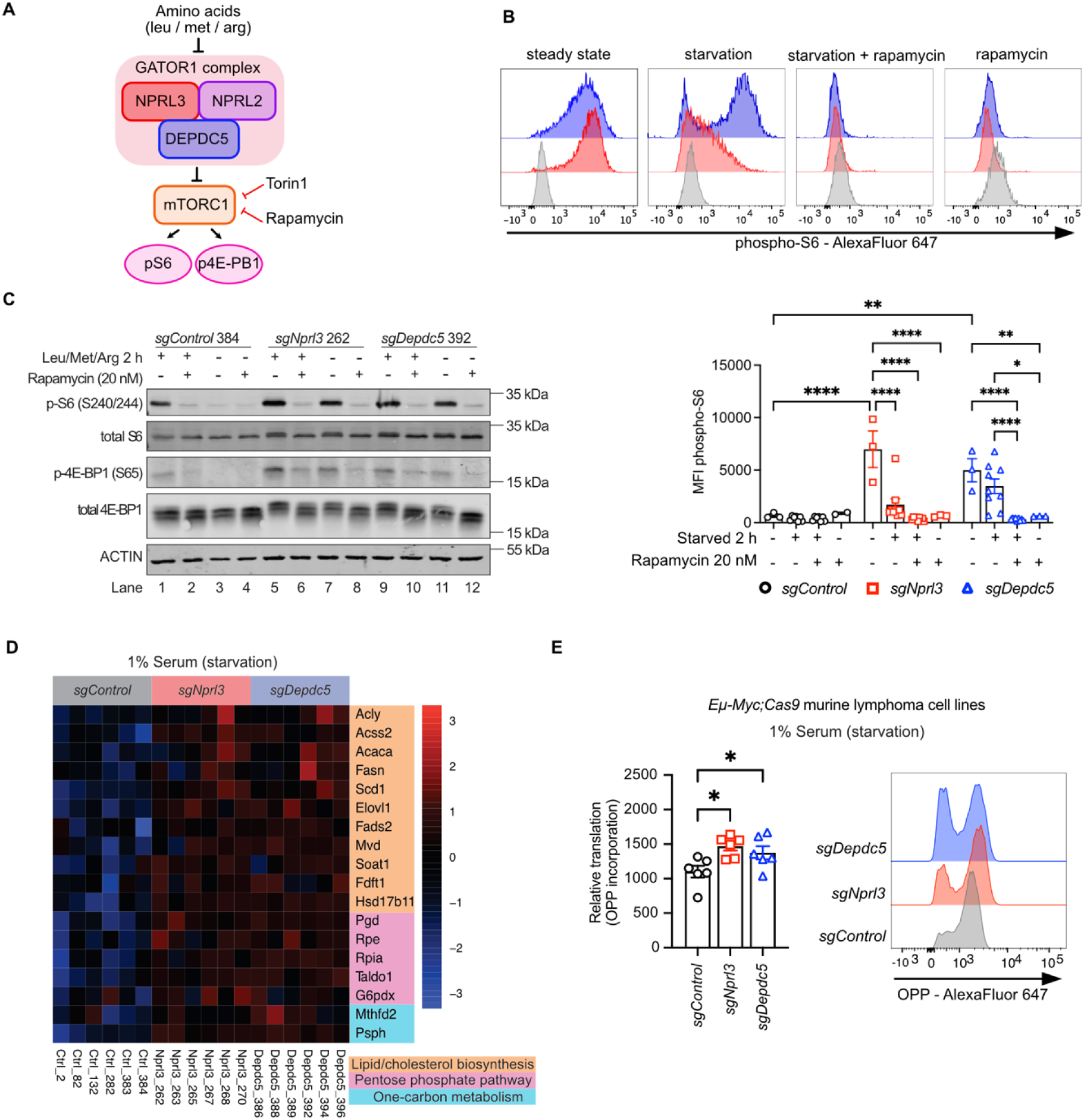
GATOR1-deficient lymphomas display alterations in mTORC1 regulated metabolic pathways. **A**, Schematic of the GATOR1 complex that negatively regulates mTORC1 signaling in response to the availability of the amino acids leucine (leu), methionine (met) and arginine (arg). mTORC1 can be directly inhibited by the drug rapamycin or the mTORC1/mTORC2 inhibitor torin1. **B**, Phospho-S6 (activated S6) protein levels (presented as mean intensity fluorescence – MFI) as measured by intracellular flow cytometry in *sgControl*, *sgNprl3* and *sgDepdc5 Eµ-Myc;Cas9* lymphoma cell lines at steady state, treatment with rapamycin (20 nM), 2 h starvation (deprived of all amino acids) and 2 h starvation plus treatment with rapamycin (20 nM). Summary graph of phospho-S6 protein levels *n* = 2-3 lymphoma cell lines per genotype across three replicate experiments are shown on the bottom and representative FACS histograms, from one representative lymphoma cell line per genotype are shown on the top. **C**, Western blot analysis of total S6, phospho-S6 (activated S6), total 4E-BP1 and phospho-4E- BP1 (activated 4E-BP1) proteins in *sgControl*, *sgNprl3* and *sgDepdc5 Eµ-Myc;Cas9* lymphoma cell lines at steady state or after 2 h starvation of the amino acids leucine (leu), methionine (met) and arginine (arg), with and without treatment with rapamycin (20 nM) for 2 h. Blotting for ACTIN served as a protein loading control. Protein sizes are indicated in kDa. **D**, The *sgNprl3* and *sgDepdc5 Eµ-Myc;Cas9* lymphoma cell lines have elevated expression of genes involved in mTORC1 regulated metabolic pathways compared to the *sgControl Eµ-Myc;Cas9* lymphoma cell lines. Heatmaps show gene expression of mTORC1 regulated metabolic pathways, lipid/cholesterol biosynthesis, pentose phosphate pathway and one-carbon metabolism from RNA sequencing analysis of *sgControl* (*n* = 6), *sgNprl3* (*n* = 6) and *sgDepdc5* (*n* = 6) *Eµ-Myc;Cas9* lymphoma cell lines after 1% serum starvation for 24 h. **E**, Relative protein translation was measured in negative control *sgControl*, *sgDepdc5* or *sgNprl3 Eµ-Myc;Cas9* lymphoma cell lines after growth for 24 h under starvation conditions (medium containing only 1% FCS) by incorporation of O-propargyl-puromycin (OPP) using Click-iT Plug Alexa Fluor 647 detected by flow cytometry. *n* = 6 independent lymphoma cell lines per genotype, 1-2 technical replicates. Ordinary one-way ANOVA statistical test was used for comparison, \**P < 0*.05, *\*\*P < 0*.005.

### GATOR1 deficiency perturbs cellular metabolism

Given that mTORC1 is a master regulator of several metabolic processes, we next explored changes in mTORC1-regulated metabolic pathways caused by loss of GATOR1 function. We performed RNA sequencing on control and NPRL3 or DEPDC5 deficient lymphoma cells following culture in either complete medium (10% fetal calf serum; FCS), or under starvation conditions (medium with only 1% FCS), the latter repressing mTORC1 activity. In both conditions, GATOR1-deficient lymphoma cells displayed elevated expression of genes involved in mTORC1 regulated metabolic pathways, relative to control cells, including lipid/cholesterol biosynthesis, pentose phosphate pathway and one-carbon metabolism (Fig. 4D, Supplemental Fig. S5A-B). This reveals that GATOR1 deficiency promotes constitutive mTORC1 activation and renders cells less sensitive to external factors that repress this pathway. No differences in mitochondrial function were seen when comparing GATOR1-deficient with control lymphoma cells (Supplemental Fig. S6A). However, consistent with known functions of mTORC1 ^22^, we observed a trend towards elevated *de novo* lipogenesis in GATOR1-deficient lymphoma cells (Supplemental Fig. S6B), as well as increased incorporation of O-propargyl-puromycin (OPP) into nascent proteins, indicative of augmented translation (Fig. 4E, Supplemental Fig. S6C). Augmented translation of the pro-survival BH3-only family member MCL-1 resulting from hyperactive mTORC1 via loss of TSC1 was reported to promote lymphomagenesis in *Eµ-Myc* transgenic mice ^32,39^. However, GATOR1 deficient lymphoma cells had comparable MCL-1 protein levels to control lymphoma cells, and MCL-1 protein expression was reduced similarly between control and GATOR1-deficient lymphoma cells following rapamycin treatment (Supplemental Figure S6D). This suggests loss of GATOR1 mediated inhibition of mTORC1 does not accelerate MYC-driven lymphomagenesis by inhibiting cell death. This conclusion was supported by GATOR1-defient lymphoma cells displaying no increased sensitivity to treatment with the MCL-1 inhibitor S63845 ^40^ (Supplemental Figure S6E). These results demonstrate that loss of GATOR1 promotes aberrant mTORC1 signaling with consequent reprogramming of cellular metabolism.

### GATOR1-deficient lymphoma cells are highly sensitive to mTORC1 inhibition

As mTORC1 is constitutively activated in GATOR1 deficient lymphomas, we hypothesized that these cells might be “addicted” to a hyperactive mTORC1 pathway, thereby potentially presenting a therapeutic vulnerability. Treatment with rapamycin (mTORC1 inhibitor) potently killed GATOR1-deficient cells *in vitro* while control *Eµ-Myc* lymphoma cells were highly resistant to this treatment (Fig. 5A). Similarly, GATOR-1 deficient lymphoma cells were sensitive to treatment with torin1, a dual mTORC1 and mTORC2 inhibitor ^41^, whereas control *Eµ-Myc* lymphoma cells were only killed by very high doses of torin1 that are not clinically relevant (Fig. 5B). Having demonstrated rapamycin sensitivity of *Eµ-Myc* lymphoma cells *in vitro*, we next assessed whether such treatment could translate into a meaningful therapeutic response *in vivo*. To this end, control or GATOR1-deficient lymphoma cell lines were transplanted into RAG1-deficient mice which were then treated with rapamycin or vehicle (control) for five consecutive days (Fig. 5C). Remarkably, most RAG1-deficient mice transplanted with GATOR1- deficient lymphomas (lacking either NPRL3 or DEPDC5) were cured by rapamycin single agent treatment, whereas control lymphomas showed no response, with all recipients presenting with severe malignant disease within 30 days (Fig. 5D). These results demonstrate that GATOR1 deficiency causes addiction of c-MYC-driven lymphoma cells to excess mTORC1 signaling, rendering them vulnerable to single agent treatment with mTORC1 inhibitors.

**Figure 5.**
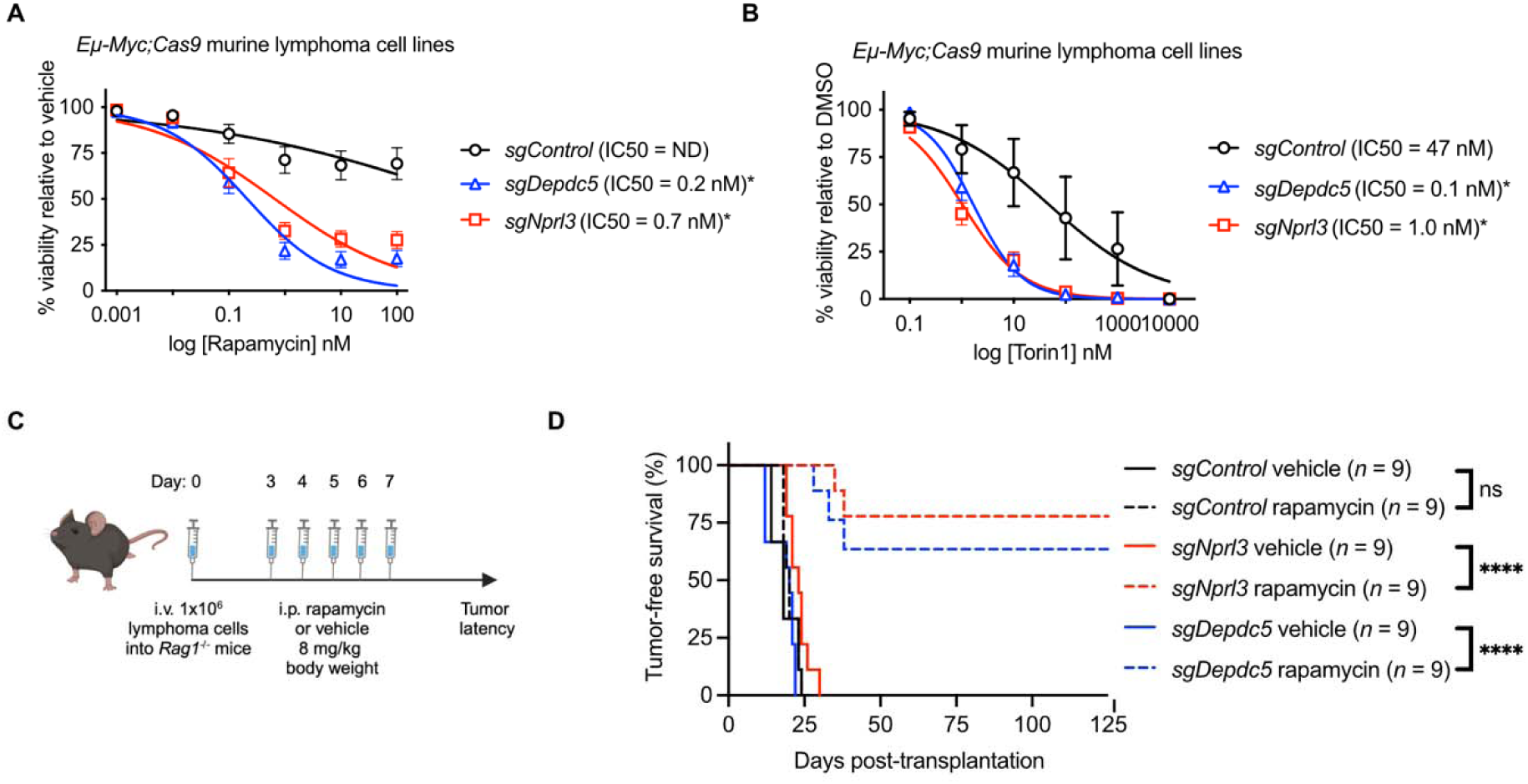
GATOR1-deficient lymphoma cells are highly sensitive to mTORC1 inhibition *in vitro* and *in vivo*. **A**, **B**, Response curves of *sgControl*, *sgNprl3* or *sgDepdc5 Eµ-Myc;Cas9* lymphoma cell lines to treatment *in vitro* with increasing doses of rapamycin **A**, or torin1 **B**. Lymphoma cell viability was measured after 24 h of treatment with drug or vehicle by staining with AnnexinV plus PI followed by flow cytometric analysis. AnnexinV/PI double negative cells were deemed viable. *n* = 3-6 lymphoma cell lines per genotype across 3 technical replicates. Data are presented as mean ± SEM, log transformed and fitting to non-linear regression. IC50 values are shown in brackets. Student *t-test* with Welch’s correction, \**P* < 0.05. **C**, Schematic of *in vivo* rapamycin treatment experiments. One million *sgControl*, *sgNprl3* or *sgDepdc5 Eµ-Myc;Cas9* lymphoma cells were transplanted i.v. into the tail vein of RAG1-deficient mice, which lack B and T cells, to prevent lymphoma rejection due to an immune response against Cas9. Two days later, mice were randomly assigned into treatment arms, receiving either rapamycin at a dose of 8 mg/kg of body weight for 5 consecutive days by i.p. injection or vehicle as a control. Mice were monitored for lymphoma growth. **D**, Tumor-free survival curve of RAG1-deficient mice that had been transplanted with 1x10^6^ negative control *sgControl*, *sgNprl3* or *sgDepdc5 Eµ-Myc;Cas9* lymphoma cell lines. *n* = 9 mice per treatment arm, with three cell lines per genotype, each injected into three recipient mice. Log-rank (Mantel-Cox) statistical test for survival curve comparison.

## Discussion

CRISPR gene knockout screens are a powerful approach to identify critical regulators of biological processes of interest ^42,43^, such as tumor suppression. While some *in vivo* CRISPR screens have been conducted in mice ^6,10,44–46^, our screen differs through employing primary non-malignant cells (HSPCs from *Eµ-Myc;Cas9* embryos) that can reconstitute the entire hematopoietic compartment and give rise to pre-B/B cell lymphomas when transplanted into lethally irradiated recipients. Maintaining coverage of a genome-wide sgRNA library *in vivo* is challenging due to several bottle neck events, such as loss of cells during transplantation or engraftment ^27^. However, our investigations demonstrate that genome-wide sgRNA library coverage is maintained *in vivo* as detected in the donor derived non-malignant hematopoietic cells present in the recipient mice. Importantly, several hits found in the lymphomas, representing known and novel candidate tumor suppressors, were identified and validated to suppress c-MYC-driven lymphomagenesis. The success of these screens indicates that this approach can be applied to other biological questions, such as identifying novel suppressors of tumorigenesis driven by different oncogenic lesions perhaps even in non-hematopoietic cell types, as long as they are amenable to sgRNA library transduction and transplantation into the relevant tissue (e.g. transduction of mammary epithelial stem cells with a sgRNA library followed by their transplantation into cleared mammary fat pads of mice).

Here, by employing the highly efficient CRISPR/Cas9 gene knockout system in primary HSPCs using an unbiased genome-wide sgRNA library screen, we reveal mTORC1 inhibitory pathway genes as potent tumor suppressor genes in c-MYC-driven lymphomagenesis. Modified cellular metabolism is a hallmark of cancer ^47^, and abnormalities in the expression of c-MYC or genes regulating various metabolic pathways are known to cooperate during cancer development ^2^. Genetic aberrations upstream of mTORC1, resulting in its constitutive activity, are frequent in many cancers. This includes activating mutations in AKT or loss of function mutations in TSC1 or TSC2, which were both hits in our screen. Notably, our screen revealed previously undescribed suppressors of MYC-driven tumorigenesis that also function in the mTORC1 inhibitory pathway, namely the GATOR1 complex subunits (NPRL3, DEPDC5 and NPRL2), with loss of either releasing negative regulation of mTORC1 and having as powerful impact on lymphoma development. Interestingly, defects in the p53 pathway are frequently selected for during lymphomagenesis in *Eµ-Myc* mice ^15,18^. However, mutations in the GATOR1 complex subunits and p53 were mutually exclusive in our murine model and in a cohort of human DLBCL. This finding is consistent with other reports where aberrant mTORC1 activation during MYC-driven lymphomagenesis through loss of TSC2 or constitutive AKT expression obviated the pressure to lose p53 pathway function ^32,39^. Together these findings suggest that genes inhibiting mTORC1 activity are potent suppressors of MYC-driven lymphoma development and obviate the pressure to lose p53 function, which is associated with poor prognosis in patients.

Interestingly, despite cancer hijacking multiple signaling pathways that converge upon mTORC1 to promote lymphomagenesis, our data suggest these may be through distinct mechanisms. Previous reports show that elevated mTORC1 signaling in the absence of TSC2 or oncogenic AKT signaling contribute to lymphomagenesis by inhibiting apoptotic cell death. Specifically, the increased mTORC1 regulation of protein translation resulted in higher levels of the pro-survival BCL-2 family member MCL-1, enabling escape from apoptotic cell death during malignant transformation ^32^. Conversely, our results suggest loss of GATOR1 inhibition of mTORC1 activity promotes lymphomagenesis through a distinct mechanism relying on enhanced global cellular metabolism rather than elevation of specific pro-survival BCL-2 proteins. This notion of distinct mechanism between GATOR1 inhibition of mTORC1 compared to aberrant activation of the PI3K/AKT/TSC signaling axis is highlighted by the profound sensitivity of GATOR1 deficient *Eµ-Myc* lymphoma cells to single agent rapamycin treatment *in vitro* and *in vivo*. In contrast, combination therapy of rapamycin with chemotherapy was required to slow disease progression in mice burdened with AKT overexpressing ^39,48^ or TSC2 deficient *Eµ-Myc* lymphomas ^32^. Notably, low levels of GATOR1 component mRNAs are predictive of poorer survival in patients with tumors expressing high levels of *c-MY*C. This indicates that this tumor suppressive process may also be critical in diverse types of human cancers. These findings identify a precision medicine opportunity for the rational and selective use of such agents in cancer therapy. Since mTOR inhibitors are already undergoing clinical trials ^49^, our work could be rapidly translated.

## Methods

### Animal husbandry and mouse models

The care and use of experimental mice were approved by and conducted according to The Walter and Eliza Hall Institute WEHI) Animal Ethics Committee guidelines. C57BL/6-Ly5.2, C57BL/6-Ly5.1 and *Rag1*/J knockout mice were obtained from the WEHI breeding facility (Kew, Victoria, Australia). *Eµ- Myc* transgenic ^3^ and constitutive *Cas9-eGFP* transgenic mice (a gift from Prof K.Rajewsky) ^50^ mice were maintained on a C57BL/6-Ly5.2 background.

### Genotyping

Mouse genotypes were determined by PCR analysis of DNA derived from earclips or embryonic tails using direct tail lysis buffer (Viagen Biotech) and Proteinase K. Mastermix was prepared with GoTaq green (Promega, M7123) containing primers (final concentration 0.5 pmol/mL) with the addition of 1 mL total DNA. PCR cycling conditions were: 94°C 4 min followed by 30 cycles (94°C for 40 sec, 55°C for 30 sec, 72°C for 60 sec) and finally 72°C for 5 min. PCR products were separated by gel electrophoresis on 2% DNA grade agarose (Bioline) gels in TAE buffer (40 mM Tris Acetate, 1 mM EDTA pH 8.0) containing ethidium bromide (0.2 mg/mL, Sigma) and imaged on GelDoc DOCTM XR+ Gel Documentation system (Bio-Rad). PCR oligonucleotide primers (Integrated DNA Technologies) and expected products were as follows:

***Eµ-Myc***: expected band size ∼900 bp

*myc-1*: 5’ -CAGCTGGCGTAATAGCGAAGAG

*myc-2*: 5’ -CTGTGACTGGTGAGTACTCAACC

***Cas9-eGFP*:** expected band size of knockin ∼112 bp and WT ∼196 bp R26F2: 5’- GCCTCCTGGCTTCTGAGGACCG

R26R2: 5’- TCTGTGGGAAGTCTTGTCCCTCC SAR: 5’-CCTGGACTACTGCGCCCTACAGA

### Transduction of HSPCs and transplantation into lethally irradiated recipient mice

Embryonic day E13.5 fetal liver cells, a rich source of HSPCs, were collected from *Eµ-Myc*;*Cas9* doubly transgenic mice (both transgenes in a heterozygous state in all experiments) on a C57BL/6-Ly5.2 background, and frozen in 90% fetal calf serum (FCS)/10% DMSO (v/v). Fetal liver cells were thawed and cultured in alpha-minimum essential medium (α-MEM) GlutaMAX (Gibco, 35050061) supplemented with 10% FCS (Gibco), 1 mM L-glutamine (Gibco, 25030149), 1 mM sodium pyruvate (Gibco, 11360070), 100 U/mL penicillin-100 µg/mL streptomycin (Gibco, 15140122), 10 mM HEPES (Gibco, 15630080), 50 µM β-mercapto-ethanol and recombinant cytokines (see below) for 48 h prior to lentiviral transduction. The cytokines IL-6 (10 ng/mL), mouse stem cell factor (100 ng/mL), thrombopoietin (50 ng/mL) and FLT-3 ligand (10 ng/mL) were kindly provided by Dr. Jian-Guo Zhang (WEHI). Lentivirus supernatant was generated as described above. Twelve-well non-tissue culture treated plates (Nunc) were coated overnight with retronectin solution (32 µg/mL in PBS, WEHI) at 4[ followed by blocking with 2% bovine serum albumin solution (Sigma-Aldrich, #A1595) in PBS at 37[ for 30 min prior to coating with viral supernatant. Viral particles were supplemented with 8 µg/mL polybrene and were centrifuged onto the retronectin coated plates at 3,500 rpm for 2 h. Supernatant was removed from the lentivirus coated plates and fetal liver cells/HSPCs were then added to wells and incubated for 24 h. Transduction efficiency was routinely ∼20-30%. Fetal liver cells transduced with vectors encoding sgRNAs were collected and pooled per sgRNA, washed in PBS, filtered through 100 µM mesh, and then injected into lethally irradiated (two doses of 5.5 Gy, 4 h apart) 7-8 week-old C57BL/6-Ly5.1 recipient mice. One fetal liver equivalent was injected into two recipient mice each (∼2-3x10^6^ cells per recipient mouse). Tumor-free survival was defined as the time from transplantation to humane ethical endpoint, which was determined by an experienced animal research technician who was blinded to the nature of the injected cells. At sacrifice, peripheral blood was collected and analyzed using an ADVIA hematology analyzer (Bayer) and hematopoietic tissues (spleen, lymph nodes, thymus, bone marrow) were collected for downstream analysis. The pooled CRISPR screens represent 6 independent cohorts. Validation experiments represent 2 independent cohorts. Each hematopoietic reconstitution experiment had internal controls of the same pool of fetal liver cells/HSPCs transduced with a negative control sgRNA (*sgControl*) to account for generational differences in the *Eµ-Myc* mouse colony and possible differences in hematopoietic reconstitution of lethally irradiated recipients between different experiments.

### Lymphoma cell transplantation and *in vivo* drug treatment

For *in vivo* drug treatment, 1x10^6^ *Eµ-Myc* lymphoma (C57BL/6-Ly5.2) cells resuspended in 200 µL sterile PBS were injected into 7-12 week-old sex-matched recipient *Rag1*^-/-^ mice by intravenous (i.v.) tail vein injection. *Rag1*^-/-^ mice were used as recipient to prevent rejection of lymphoma cells owing to an immune response to Cas9. After three days, mice were treated for five consecutive days with vehicle or 8 mg/kg body weight rapamycin (LC laboratories) by intra-peritoneal (i.p.) injection. Rapamycin was first dissolved in 100% ethanol, stored at -20[ and further diluted in vehicle, 5.2% PEG 400 and 5.2% Tween 80, immediately before use. Mouse survival time was defined as the time from lymphoma cell line injection until mice were sick and had to be euthanized according to our animal ethics guidelines. This was judged by an experienced animal technician who was blinded to the nature of the lymphoma cells that had been injected and the treatment of the mice.

### Lentiviral constructs and sgRNAs

A positive control sgRNA targeting *p53* (5’-GGCAACTATGGCTTCCACCT) and a negative control sgRNA targeting human-*BIM* (5’-GCCCAAGAGTTGCGGCGTAT) were derived from the constitutive sgRNA FUGW expression vector previously described ^13^. The genome-wide mouse lentiviral CRISPR sgRNA library used for *in vivo* screening experiments, YUSA v1 was a gift from Kosuke Yusa (Addgene #50947) ^23^. Two independent sgRNAs for *in vivo* validation experiments of hits and a negative control sgRNA targeting human *NLRC5,* (Supplemental Table 1) were obtained from the Merck CRISPR glycerol stock arrayed mouse sgRNA library available at WEHI ( Sigma Aldrich #<u>MSANGERG</u>).

### Lentivirus production

Lentiviruses were produced by transient transfection of HEK293T cells in 10 cm^2^ dishes with 10 µg of vector DNA and packaging lentiviral constructs pMDL (5 µg), RSV (2.5 µg), and pENV (5 µg) by the standard calcium phosphate precipitation method previously described ^51^. Viral supernatants were collected 48-72 h after transfection and passed through a 0.45 mM filter prior to transduction of cells.

### Next generation sequencing

To identify pooled CRISPR sgRNAs, genomic DNA was isolated from spleens of reconstituted lymphoma bearing mice (containing malignant and normal hematopoietic cells) using DNeasy Blood and Tissue Kit (Qiagen) according to the manufacturer’s instructions. Integrated sgRNAs in each lymphoma sample were amplified from 100 ng of genomic DNA using GoTaq Green Master Mix (Promega) and indexing primers with unique overhangs ^24^. Amplicons were pooled, cleaned up using Ampure XP beads (Beckman Coulter) and sequenced on Illumina Nextseq or Miseq platform. Enriched sgRNAs per lymphoma were calculated by number of reads mapping to each sgRNA in the library as a proportion of the total number of reads within this lymphoma sample. To identify InDels or mutations in individual target genes, DNA was extracted from lymphoma derived cell lines as described above and PCR amplified using sgRNA specific primers (Supplemental Table 2, *p53* primers as previously described ^25^ and GoTaq Green Master Mix (Promega). PCR products were amplified a second time using indexing primers, pooled, purified using Ampure XP beads (Beckman Coulter) and sequenced on an Illumina Miseq platform. Reads were aligned to WT reference sequences obtained from Ensembl.

### Cell culture

HEK293T cells were cultured in DMEM medium supplemented with 10% FCS. Prior to virus production, HEK293Ts were cultured in DME Glutamax medium (Gibco) supplemented with 10% FCS and 25 mM HEPES. *Eµ-Myc* lymphoma cell lines derived from tumors of sick mice were maintained in FMA medium as previously described ^52^. All cell lines were regularly verified as mycoplasma negative (MycoAlert mycoplasma detection kit; Lonza, LT07-118) and authenticated by STR profiling at the Australian Genome Research Facility.

### *In vitro* drug treatment assays

To assess drug sensitivity of *Eµ-Myc* lymphoma cells, 5 x 10^4^ cells were plated in triplicate into 96-well flat-bottom plates. Cells were treated with rapamycin (Cell Signaling Technology, 9904), torin1 (Cell Signaling Technology, 14379), nutlin-3a (Cayman Chemical, 18585), etoposide (Ebewe Pharmaceuticals Ltd, A-4866) or the MCL-1 inhibitor S63845 (Active Biochem, 6044) at the indicated concentrations. Cell viability was assessed after 24 h of treatment by staining cells with 2 µg/mL propidium iodide (PI) and AnnexinV conjugated to Alexa Fluor 647 (produced in-house) followed by flow cytometric analysis using a Fortessax20 flow cytometer (Becton Dickinson), 10,000 events were recorded per sample. Data were analyzed using FlowJo analysis software. Cell viability was normalized to control cells treated with an equivalent volume of vehicle, either dimethyl sulfide (DMSO), water or 100% ethanol.

### Flow cytometric analysis

Immunophenotyping of lymphoma cells was performed on single cell suspensions of primary lymphoma tissue from sick mice or *Eµ-Myc;Cas9* lymphoma derived cell lines stained with fluorochrome-conjugated antibodies against B220 (RA3-6B2), IgM (5.1), IgD (11-26C), T cell antigen receptor beta (TCRβ, H57-597) and CD19 (ID3) in PBS supplemented with 5% FCS and Fcγ receptor block (2.4G2 hybridoma supernatant, WEHI). Donor derived cells were identified as CAS9, i.e. eGFP positive and sgRNA, i.e. CFP or BFP positive. Propidium iodide (PI), final concentration of 2 mg/mL, was used for excluding dead cells during analysis on a Fortessax20 flow cytometer (Becton Dickinson).

### Intracellular flow cytometric analysis for phosphorylated S6

To assess the levels of phosphorylated (i.e. activated) S6 by flow cytometry, lymphoma cells were washed in PBS then resuspended in either FMA medium or Hanks Balanced Salt Solution (HBSS), for starvation condition, without or with supplementation with 20 nM rapamycin and incubated for 2 h at 37°C in 10% CO_2_. Cells were spun down and 1x10^5^ cells were harvested, washed in PBS and stained with green fixable viability dye (Thermo Fisher Scientific). Cells were fixed for 1 h using the eBioscience^TM^ Foxp3/Transcription Factor Staining Buffer Set (Thermo Fisher Scientific) and stained according to the manufacturer’s instructions. The antibodies used were directed against phospho-S6 (i.e. activated S6) conjugated to Alexa Fluor 647 (Cell Signaling Technology, 4851). Cells were then analyzed on an LSR IIC flow cytometer (Becton Dickinson). Data were analyzed using FlowJo analysis software and are presented as geometric mean fluorescence intensity (MFI).

### Western blotting

Total protein extracts were prepared from *Eµ-Myc;Cas9* lymphoma cell lines and primary lymphomas containing the indicated sgRNAs by lysis in RIPA buffer (50 mM Tris-HCl, 150 mM NaCl, 1% NP-40, 0.5% DOC, 0.1% SDS) containing complete protease inhibitor cocktail (Roche) and phosphatase inhibitor cocktail (PhosphoSTOP^TM^, Roche). Protein concentration was quantitated by using the Bradford assay (Bio-Rad). Between 10 to 30 µg of protein were prepared in Laemmli buffer boiled for 5 min and proteins size separated by SDS-PAGE using NuPAGE 10% Bis-Tris 1.5 mm gels (Life Technologies). Proteins were then transferred onto nitrocellulose membranes (Life Technologies) using the iBlot membrane transfer system (Thermo Fisher Scientific). Membranes were incubated with the primary antibodies and then the HRP-conjugated secondary antibodies detailed in the Supplemental Table 3, diluted in PBS containing 10% bovine serum albumin with 0.1% Tween20. Luminata Forte Western horseradish peroxidase (HRP) substrate (Merck Millipore) was used for developing the signal and membranes were imaged and analyzed using the ChemiDoc Imaging System with ImageLab software (Bio-Rad). Cell lysates from the p53 mutant *Eµ-Myc* lymphoma cell line (EMRK1172) ^53^ served as a positive control and the p53 wild-type *Eµ-Myc* lymphoma cell line (EMRK1184) ^54^ served as a negative control for high levels of mutant p53 protein and p19^ARF^ protein. For Western blot detection of phosphorylated proteins, lymphoma cell lines were cultured for 2 h in FMA medium supplemented with 10% dialyzed heat-inactivated FBS, or FMA medium lacking the amino acids leucine, methionine and arginine, supplemented with 10% dialyzed heat-inactivated FBS. Concurrently, these cells were treated with DMSO (vehicle) or 20 nM rapamycin (Cayman Chemical, 20022). Cells were washed with PBS and lysed in ice-cold RIPA buffer containing a protease inhibitor cocktail (Sigma-Aldrich, P8340) and phosphatase inhibitors (PhosSTOP, Roche, 4906837001). Lysates were resolved by SDS-PAGE and transferred onto a nitrocellulose membrane (see above) followed by immunoblotting. Membranes were probed with primary antibodies (see Supplemental Table 3) followed by IRDye secondary antibodies (LI-COR Biosciences, goat anti-rabbit IgG,1:20000, 926-32211 and goat anti-mouse IgG, 1:50000, 926- 68070). Membranes were developed using infrared imaging (LI-COR Odyssey CLx).

### RNA sequencing

Lymphoma cells were cultured in FMA medium containing full serum (10% FCS) or under starvation conditions (FMA medium containing only 1% FCS) for 24 h prior to harvesting. RNA was isolated from *Eµ-Myc* lymphoma cells using a NucleoSpin RNA kit (Macherey-Nagel) following the manufacturer’s instructions. The 3’ libraries were sequenced with an Illumina NextSeq 500, with single-end 75 bp reads to a depth of 15 M reads per sample. FastQ files were trimmed using Cutadapt (v1.16) and mapped to the mouse reference genome (mm10 GRCm38) using HISAT2 (v2.1.0). Samtools (v1.4.1) was used to convert SAM files to BAM format and to sort BAM files. Counts were generated using featureCounts (v1.6.0). Filtering and normalization of data were carried out using edgeR (v3.16) ^55^ and differential gene expression analysis was carried out using limma-voom (v3.4.6) ^56^. Heatmaps were generated using pheatmap (v1.0.12).

### Lipogenesis assay

A previously described protocol, with minor modifications, was employed ^57^. Briefly, *Eµ-Myc* lymphoma cells grown in 6-well plates were spiked with 0.2 μCi/mL [1-^14^C]-acetate (PerkinElmer; NEC084H001MC) for the final 4 h of the experimental period. Cells were pelleted and washed twice with PBS, before lysis in 150 μL 0.5% Triton X-100. Lipids were extracted with 500 μL 2:1 (v/v) chloroform/methanol followed by centrifugation at 1000 rpm for 20 min. 150 μL ^14^C-labelled lipids from the denser organic fraction were combined with 4 mL OptiPhase HiSafe 3 liquid scintillation cocktail (PerkinElmer; 1200.437) and radio-labeling was quantified using a Tri-Carb 2910 TR Liquid Scintillation Analyzer (PerkinElmer). ^14^C-acetate incorporation was normalized to cell number.

### Seahorse mito stress test

Three of each *sgControl, sgNprl3* and *sgDepdc5 Eµ-Myc;Cas9* lymphoma cell lines were treated with 0.45 nM of rapamycin or vehicle (DMSO) for 3 h. Drug was then washed out and 1x10^5^ cells plated in 5 replicate wells for immediate analysis using the Mito Stress Test Kit (Agilent, 103015-100) according to the manufacturer’s instructions and using the following drug concentrations: 1.5 µM oligomycin, 2.0 µM FCCP, 0.5 µM Rotenone/Antimycin A. Mitochondrial function was analyzed on the Seahorse XFe96 (Agilent) analyzer. Plots were prepared using GraphPad Prism software.

### Puromycin incorporation assay to determine rate of translation

A previously described protocol, with minor modifications, was employed ^58^. *Eµ-Myc* lymphoma cells grown in 6-well plates were spiked with 20 µM O-propargyl-puromycin (Thermo Fisher Scientific, C10459) for the final 30 min of the experimental period. Cells were pelleted and washed with PBS. Cells were then fixed with 3.7% paraformaldehyde for 20 min, followed by a 20 min permeabilization with 0.5% Triton-X. Cells were stained using the Click-iTTM Plus Alexa FluorTM 647 Picolyl Azide Toolkit (Thermo Fisher Scientific, C10643) according to the manufacturer’s instructions. Puromycin incorporation was analyzed using a LSR FortessaTM X-20 flow cytometer (BD Biosciences).

### Mutational co-occurrence analysis in human DLBCL

Data pertaining to mutations in *p53*, *DEPDC5* and *NPRL2* in human DLBCL samples were obtained from ^59–61^ and the TCGA database (https://www.cancer.gov/tcga). Waterfall plots were generated for the samples (*n* = 73) using the waterfall function of the GenVisR ^62^ R package. Fisher’s exact test was used to test for significance of co-occurrence of mutations.

### Human Burkitt lymphoma analysis

Gene expression profiles and clinical characteristics were obtained from Lacy et al ^63^. Human Burkitt lymphoma samples (*n* = 83) were selected for patient survival analysis. The gene expression levels for *NPRL2*, *NPRL3* and *DEPDC5* across samples were divided into quartiles. Samples were divided into quartiles by *GATOR1-*low group, defined as patients with *NPRL2*, *NPRL3*, or *DEPDC5* expression in the bottom quartile. Otherwise, the sample was designated as *GATOR1*-high. The Kaplan-Meier method was used to estimate the survival functions among patient groups, and the log-rank test was used to test the differences in the overall survival between the selected groups computed using the survminer R package (v0.4.9).

### Human TCGA pan-cancer analysis

TCGA pan-cancer RSEM ^64^ batch normalized mRNA expression data for cancers from patients with expression values for *MYC*, *NPRL2*, *DEPDC5* and *NPRL3* was downloaded from the cBioPortal (*n* = 10 967 samples) ^65,66^ on July 30^th^, 2021. The R statistical software (v4.1.2) was used to conduct patient survival analysis. The Kaplan-Meier survival statistics were computed using R package ‘survival’ (v3.2- 13) and plotted using R package ‘survminer’ (v0.4.9). Clinical data relating to tumor entity or treatment grade were not available across all cancers. Therefore, further analysis was conducted with all factors present across all samples. Patient samples were first stratified on *MYC*-high expression, defined as samples where *MYC* mRNA expression was higher than the median expression of all samples (*n* = 5035). Then samples were divided by GATOR1 component mRNA expression into quartiles and patient survival was compared between the *GATOR1-low* group defined as tumors with *NPRL2* or *DEPDC5* or *NPRL3* mRNA expression in the bottom quartile versus the *GATOR1-high* group defined as *NPRL2* and *DEPDC5* and *NPRL3* mRNA expression in the top quartile.

## Statistical analysis

GraphPad Prism software (v8) was used to plot Kaplan-Meier mouse survival curves. For comparing two groups, Student’s *t*-test was used, and for comparing more than two groups multiple t-test or two-way ANOVA were performed, unless otherwise stated in the figure legends. Data are presented as mean ± standard error of the mean (SEM), unless otherwise stated in figure legends. Significance was deemed where *P* value is less than 0.05, ns = no significant difference. For the exploratory analysis of the sgRNA counts, heatmaps and pie charts were plotted using R package ‘ggplot’ (v2_3.3.5), ‘o’ represents the number of observations of sgRNAs or genes.

## Acknowledgments

The authors thank all members of the Blood Cells and Blood Cancer Division at The Walter and Eliza Hall Institute (WEHI) and the Genome Engineering and Cancer Modelling Program at the Olivia Newton-John Cancer Research Institute (ONJCRI) for their support, advice, and sharing of reagents; G Siciliano and other Bioservices staff at WEHI for technical assistance with the *in vivo* experiments and animal husbandry; A Aeslop and the Screening lab at WEHI for the glycerol stock sgRNAs for target validation. We thank B Haley and K Potts for their constructive feedback on the manuscript. Visual schematics were created using BioRender.com. This work was supported by grants and fellowships from the Australian National Health and Medical Research Council (NHMRC) (Project Grants 1159658,1186575 and 1145728 to MJH, 1143105 to MJH and AS, Ideas Grants 2002618 and 2001201 to GLK, 2004212 and 2012313 to KKB, Program Grant 1113133 to AS and Fellowships 1020363 to AS, 2017971 to MJH, 1102742 to JEV), the Leukemia and Lymphoma Society of America (LLS SCOR 7015-18 to AS, GLK, MJH), the Cancer Council of Victoria (project grant 1147328 and 2021 Grant In Aid to MJH, 1052309 to AS ,1147328 to GLK and Venture Grant to MJH and AS), Victorian Cancer Agency (MCRF Fellowship 17028 to GLK and 17020 to KKB), Phenomics Australia (to AJK and MJH) the estate of Anthony (Toni) Redstone OAM (AS and GLK), the Craig Perkins Cancer Research Foundation (GLK), the Dyson Bequest (GLK), the Harry Secomb Trust (GLK), the University of Melbourne Research Training scholarship (MAP) and the Uehara Memorial Foundation and JSPS Grant-in-Aid for Research Activity Start-up 20K22854 (SM). This work was made possible by operational infrastructure grants through the Australian Government Independent Research Institute Infrastructure Support Scheme (361646 and 9000220) and the Victorian State Government Operational Infrastructure Support Program.

## Author contributions

M.J.H. and A.S. conceived and designed the study; M.A.P., S.M., Y.D., G.H., Z.W., and C.K. conducted experiments; S.V., K.E.T. and K.K.B. conducted metabolomic studies and relating analyses; G.G. and

A.T.P. conducted pan-cancer analysis and CRISPR screen library analyses; S.S. and A.D.J. conducted survival analyses on human Burkitt lymphoma samples and p53 mutual exclusivity analysis in human DLBCL; S.W. ran NGS; L.W. provided technical mouse support; G.P., A.W., D.K. and S.D. provided reagents; M.A.P., S.M., Y.D., S.V., K.E.T., K.K.B., A.S. and M.J.H. analyzed and interpreted results; M.A.P., E.J.L., M.J.H. and A.S. drafted and edited the manuscript with feedback from all authors.

