## Supplementary Information and Figures for "Genome-wide *in vivo* CRISPR screens identify GATOR1 as a potent tumor suppressor in MYC-driven lymphomagenesis"

**Supplemental Figure S1**

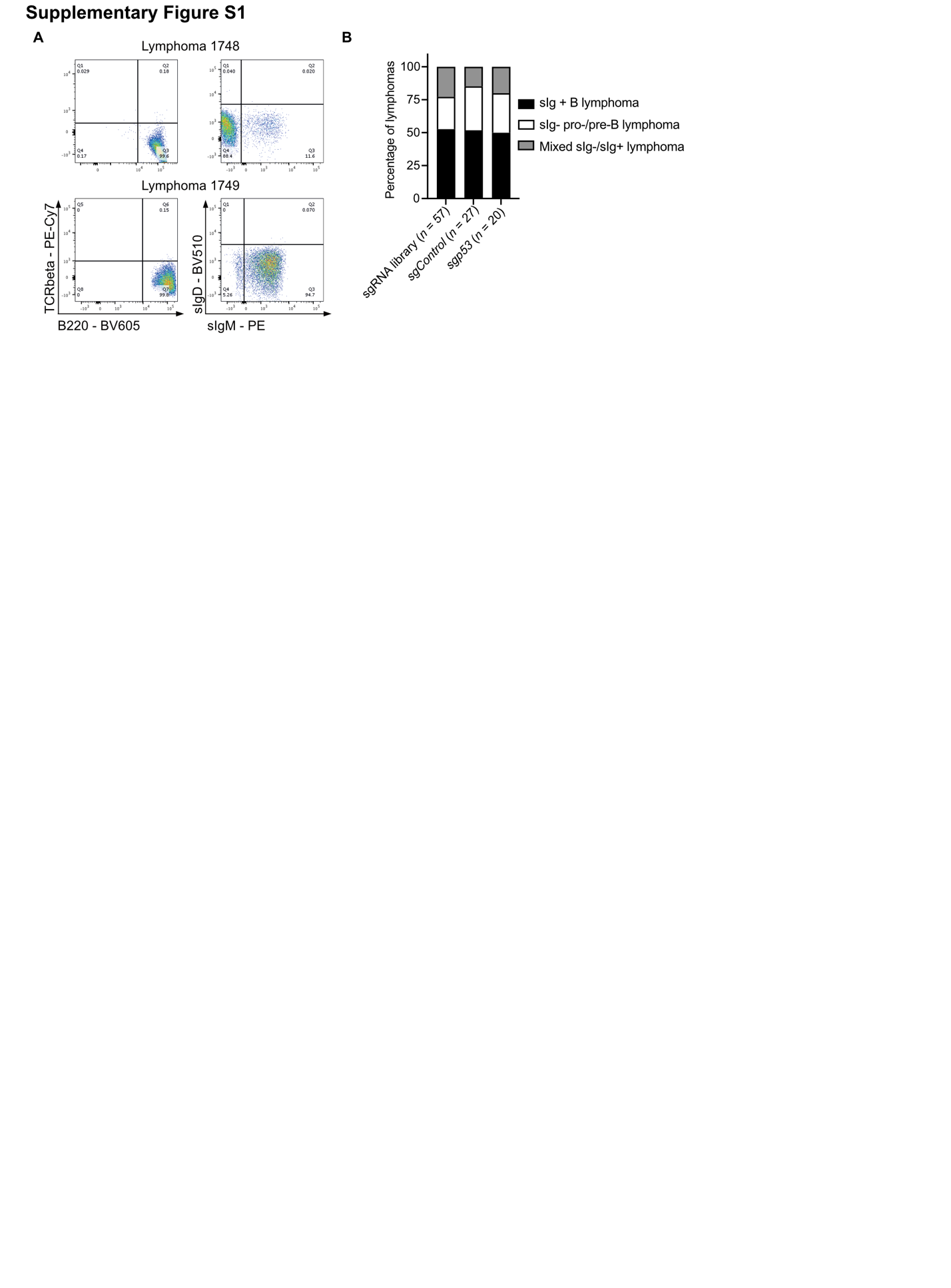
**Supplemental Figure S1**

Immunophenotyping of lymphomas from mice that had been transplanted with *Eµ-Myc;Cas9* HSPCs that had been transduced with a negative control sgRNA (*sgControl*), a *sgp53* (positive control) or the whole-genome sgRNA library HSPCs. **A**, Representative flow cytometry plots of lymphoma immunophenotyping. **B**, Proportions of B220+ sIg- pro-B/pre-B cell, B220+ sIg+ B cell, or ‘mixed’ pre-B cell/B cell (if sIg- and sIg+ lymphoma cells were both present). n represents total numbers of lymphomas analyzed per group indicated in brackets below each bar.

**Supplemental Figure S2**

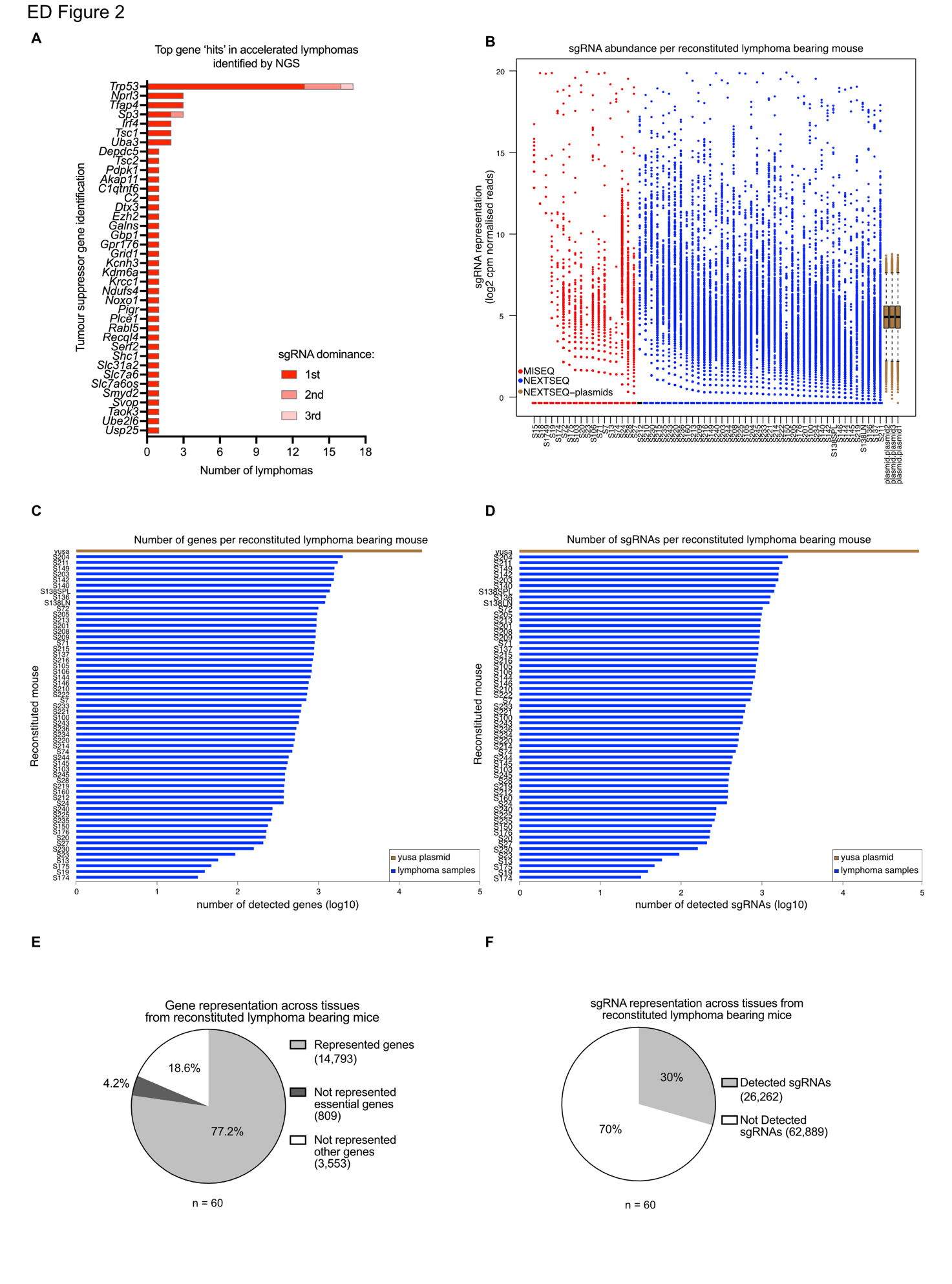

**Supplemental Figure S2**

DNA sequencing analysis from lymphoma burdened spleens (also containing sgRNAs present in non-malignant hematopoietic cells) from all mice that had been transplanted with HSPCs that had been transduced with the whole genome sgRNA that were deemed to have shown accelerated lymphoma development (*n* = 60/113). **A**, Tumor suppressor gene identification from mice with accelerated lymphoma development ranked by the dominance of its sgRNA as a proportion of total sequencing reads within a lymphoma sample, and the number of lymphomas a sgRNA for its target gene was detected as the first, second or third most dominant sgRNA. **B**, Abundance of sgRNAs detected in each lymphoma sample measured as log2 counts per million (cpm) normalized reads. Lymphoma samples colored by sequencing platform Miseq (red) or Nextseq (blue). Triplicate sequencing of YUSA library plasmid stock (brown boxplots) for comparison. **C**, Number of genes represented by detection of at least one sgRNA in the YUSA library plasmid stock and each individual lymphoma sample sequenced. **D**, Number of sgRNAs detected in the YUSA library plasmid stock and each individual lymphoma sample sequenced. **E**, Of the 19,150 genes targeted by the genome-wide sgRNA library, 77.2% (14,793 genes) were represented by >1 sgRNA, 4.2% (809 genes) were not represented by any sgRNA and are classified as essential genes and 18.5% (3,553 genes) were not represented by any sgRNA but are not essential. **F**, Of the sgRNAs in the genome-wide library, ~30% (26,262) were detected and ~70% (62,262) were not detected.

**Supplemental Figure S3**

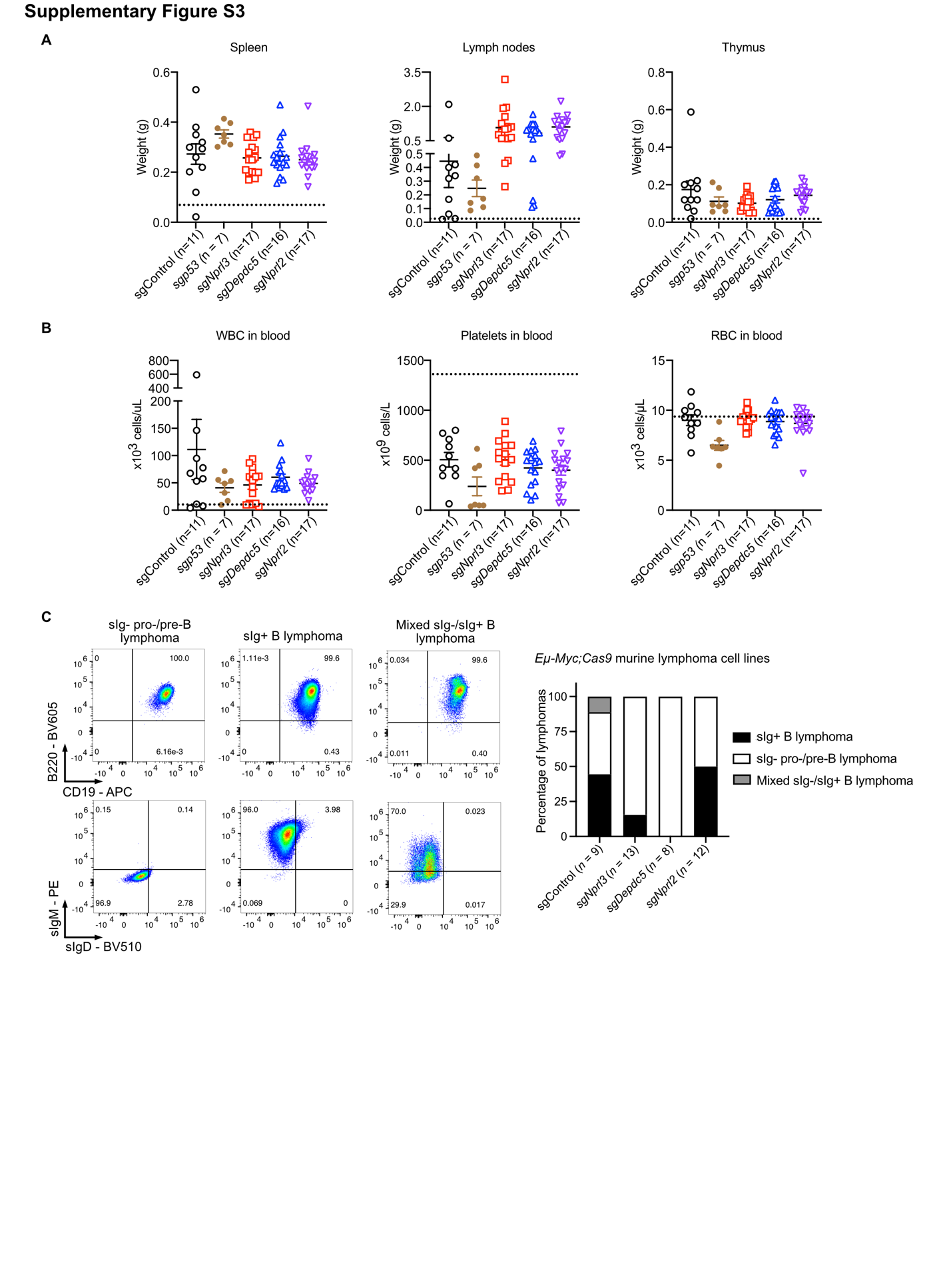

**Supplemental Figure S3**

**A**, Weights of the spleen, lymph nodes (combined axillary, brachial, inguinal) at time of sacrifice of sick recipient mice that had been transplanted with *Eµ-Myc;Cas9* HSPCs that had been transduced with the indicated sgRNAs. **B**, Enumeration of total white blood cells (WBC), platelets and red blood cells (RBC) in peripheral blood of the mice mentioned in (**A**) by ADVIA analysis. Each dot represents one mouse. Data are presented as mean ± SEM. The dotted black line represents the lymphoid tissue weights and blood cell counts averaged from six healthy six-week-old C57BL/6 mice. **C**, Representative flow cytometry plots (left) and summary graph (right) of immunophenotyping for classification of lymphoma cell lines derived from lymphomas of recipient mice that had been transplanted with *sgControl, sgNrpl3, sgDepdc5* or *sgNprl2* transduced *Eµ-Myc;Cas9* HSPCs. Classifications are: B220+ sIg- (pro-B/pre-B cell lymphoma), B220+ sIg+ (B cell lymphoma), or ‘mixed’ (pre-B/B cell lymphoma) if both sIg- and sIg+ lymphoma cells were present. *n* = number of lymphoma cell lines analyzed per sgRNA cohort indicated in brackets.

**Supplemental Figure S4**

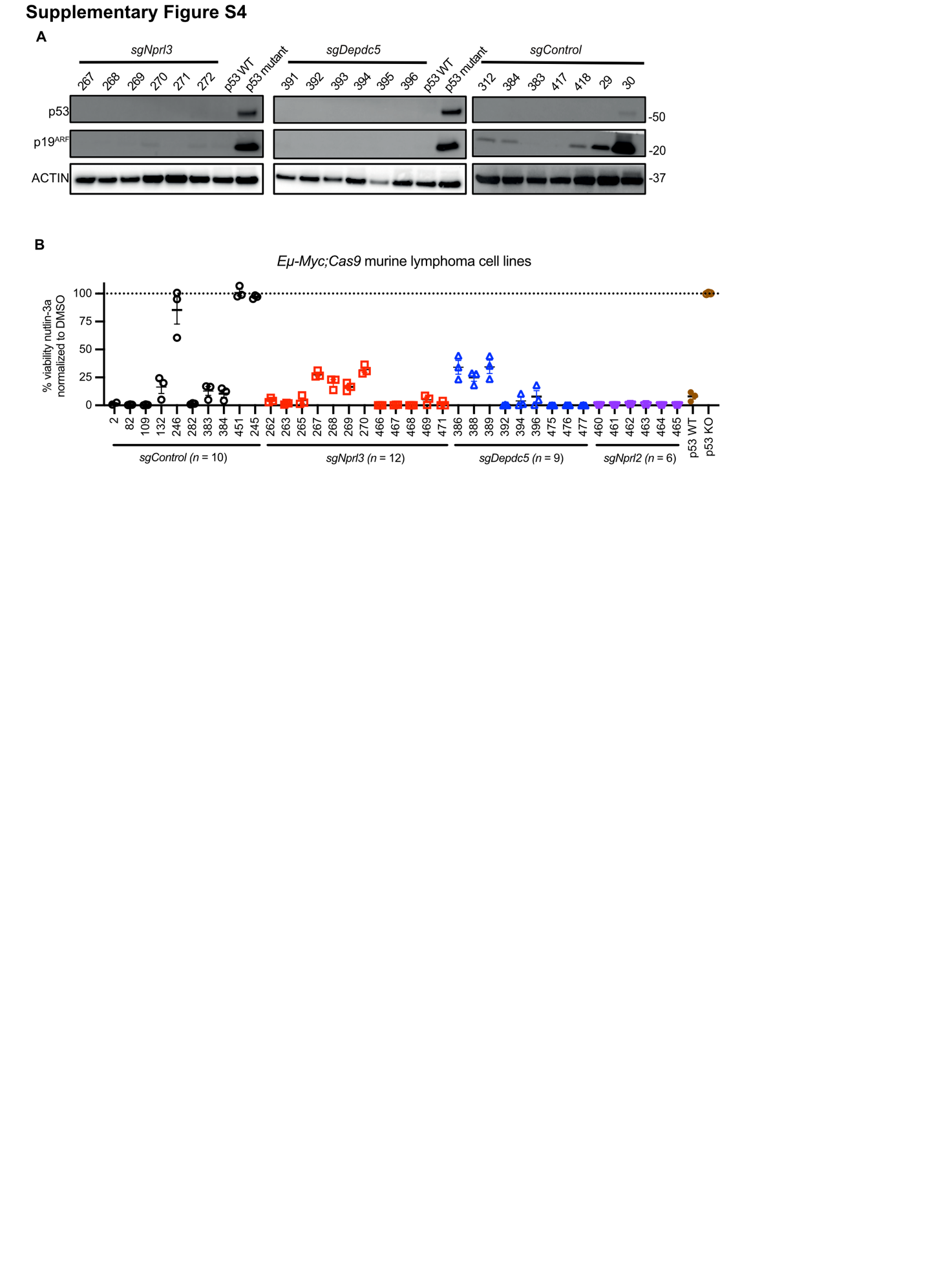

**Supplemental Figure S4**

**A**, Representative Western blot analysis relating to summary graph in Fig. 3A of p53 and p19^ARF^ protein levels to assess p53 functional status in primary *sgControl*, *sgDepdc5* or *sgNprl3* *Eµ-Myc;Cas9* lymphoma samples. Probing for ACTIN served as a protein loading control. Protein size standards are indicated in kDa. Cell lysates from *p53* WT (EMRK1184)^1^ and *p53* mutant (EMRK1172) *Eµ-Myc* lymphoma cells ^2^ served as negative and positive controls, respectively, for high levels of mutant p53 and p19^ARF^ proteins. **B**, Testing of p53 function by determining sensitivity of lymphoma cells to the MDM2 inhibitor nutlin-3a relating to summary graph in Fig. 3B. Percentage cell survival measured by flow cytometry (live cells were identified as Annexin V/PI double negative) of independent *sgControl*, *sgDepdc5*, *sgNrpl3* or *sgNprl2* *Eµ-Myc;Cas9* lymphoma cell lines after 24 h of treatment with 10 µM nutlin-3a, normalized to DMSO (vehicle control) treatment.

**Supplemental Figure S5**

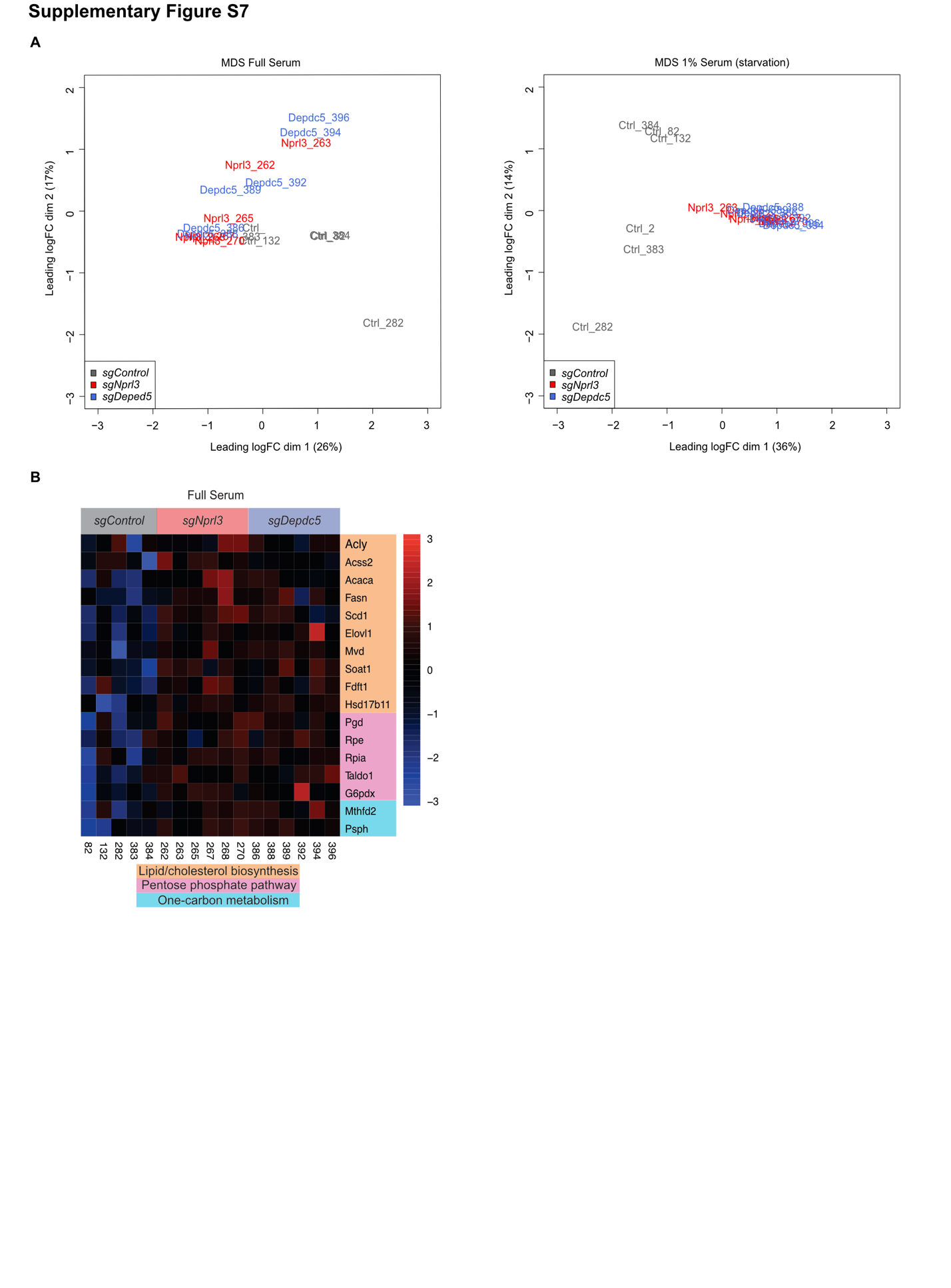

**Supplemental Figure S5**

RNA sequencing data. **A**, MDS plots showing clustering of *sgControl*, *sgNprl3* and *sgDepdc5* *Eµ-Myc;Cas9* lymphoma cell lines grown either in complete serum containing medium (10% FCS, left) or in starvation conditions (FMA medium containing only 1% FCS, right). **B**, *sgNprl3* and *sgDepdc5* *Eµ-Myc;Cas9* lymphoma cell lines have elevated expression of genes involved in mTORC1 regulated metabolic pathways compared to *sgControl* *Eµ-Myc;Cas9* lymphoma cell lines. Heatmaps show gene expression of mTORC1 regulated metabolic pathways, lipid/cholesterol biosynthesis, pentose phosphate pathway and one-carbon metabolism from RNA sequencing analysis of *sgControl* (*n* = 5), *sgNprl3* (*n* = 6) and *sgDepdc5* (*n* = 6) *Eµ-Myc;Cas9* lymphoma cell lines grown in complete medium containing 10% FCS.

**Supplemental Figure S6**

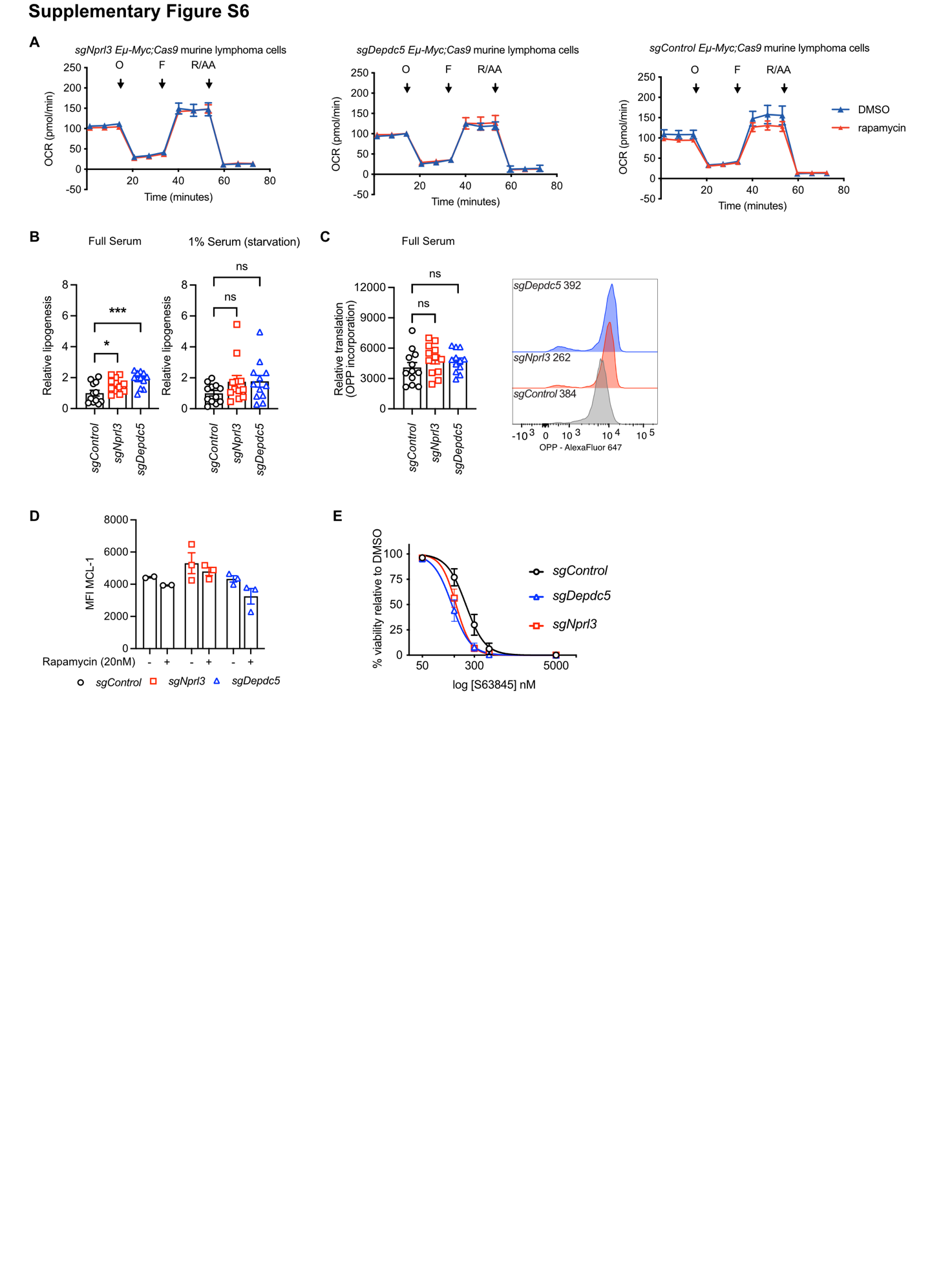

**Supplemental Figure S6**

**A**, Oxygen consumption rate (OCR) measured by using the Seahorse XFe mito stress test assay in *sgControl*, *sgDepdc5* or *sgNprl3* *Eµ-Myc;Cas9* lymphoma cell lines that had been treated for 3 h with 0.45 nM rapamycin (~IC50 dose). Representative graph of data from one lymphoma cell line per genotype where *n* = 3 lymphoma cell lines per genotype with 5 technical replicates per condition. **B**, *De novo* lipogenesis measured through incorporation of ^14^C-acetate at 24 h in *sgControl*, *sgDepdc5* or *sgNprl3 Eµ-Myc;Cas9* lymphoma cell lines when growing in full serum containing medium (10% FCS) or in 1% serum containing medium (starvation condition) for 24 h. **C**, Protein translation was measured in *sgControl*, *sgDepdc5* or *sgNprl3 Eµ-Myc;Cas9* lymphoma cell lines when growing in full serum containing medium (10% FCS) for 24 h by incorporation of O-propargyl-puromycin (OPP) by using Click-iT Plug Alexa Fluor 647 that was detected by flow cytometry. *n* = 6 lymphoma cell lines per genotype, 1-2 technical replicates. Ordinary one-way ANOVA statistical test was used for comparison, **P < 0*.05, ***P < 0*.005. **D**, MCL-1 protein levels summary graph (presented as mean intensity fluorescence – MFI) as measured by intracellular flow cytometry in *sgControl*, *sgNprl3* and *sgDepdc5* *Eµ-Myc;Cas9* lymphoma cell lines at steady state with or without rapamycin (20 nM). *n* = 2-3 lymphoma cell lines per genotype across two replicate experiments. **E**, Response curves of *sgControl*, *sgNprl3* or *sgDepdc5 Eµ-Myc;Cas9* lymphoma cell lines following treatment with increasing doses of etoposide or the MCL-1 inhibitor S63845. Cell viability was measured after 24 h of treatment with drug or vehicle by staining with AnnexinV plus PI followed by flow cytometric analysis. AnnexinV/PI double negative cells were deemed viable. *n* = 3-6 lymphoma cell lines per genotype across 3 technical replicates. Data are presented as mean ± SEM, log transformed and fitting to non-linear regression.

**Supplemental Materials and Methods**

**Potts, Mizutani, Deng, et al.,**

**Supplemental Table 1:** Merck glycerol stock sgRNA sequences for genes of interest

| **Target gene sgRNA** | **Sequence** |
| --- | --- |
| *sgNLRC5* | 5’- GCTGCAGAAGTGTCAGCTCCAGG |
| *sgNprl3 #1*  *sgNprl3 #2* | 5’ - TGACAGCATATCTGCTCCGTGG  5’ - GTTATTCTGGCAACAATTTTGG |
| *sgDepdc5 #1*  *sgDepdc5 #2* | 5’ - TTGCAAGTCAAGTCGCTTAAGG  5’ - TCACATAGACATCTTGATAAGG |
| s*gNprl2 #1*  *sgNprl2 #2* | 5’ - GAACTGTTTGACACGGTCC  5’ - GGCTTTGTGTGTGACGCTC |

**Supplemental Table 2:** Target gene specific NGS primers

| **Target gene sgRNA** | **Target site primers** |
| --- | --- |
| *Nprl3 #1* | FWD 5’ - AAGCAGTGGGTTTTAATAGCT  REV 5’ - CAGTCCCAGATGTGCACTTCC |
| *Nprl3 #2* | FWD 5’ - TGTGGGTACCTGGGCATGGTG  REV 5’ - CACATTCACC AGCATCAACAT |
| *Depdc5 #1* | FWD 5’ - GTTTACTTTTTAACTGAGGTA  REV 5’ - GCATCAACTATTGCTCCTGGGGT |
| *Depdc5 #2* | FWD 5’ - TTAACTGATTCATATTTTTTT  REV 5’ - CTGAGGACTTAAAAGCAGCC |
| *Nprl2 #1* | FWD 5’ - GGAGCTCTCTCTCATGGCTG  REV 5’ - AGCCCAGGTTGAAGAGGAGA |
| *Nprl2 #2* | FWD 5’ - AAGAAGCTGATTGGCTGCCC  REV 5’ - CAGCACAGTGAGTGACCACA |

**Supplemental Table 3:** Antibodies used for Western blot analysis

| **Antibody** | **Clone** | **Source** | **Dilution** |
| --- | --- | --- | --- |
| **Primary antibodies** | | | |
| mouse anti-HSP70 | N6 | Dr. R. Anderson, Olivia Newton-John Cancer Research Institute, Melbourne Australia | 1:1000 |
| mouse anti-p53 | CM5 | Novocastra^TM^ Leica Biosystems,  NCL-L-p53-CM5p | 1:2000 |
| mouse anti-P19/ARF | 5.C3.1 | Rockland, 200-501-891 | 1:1000 |
| rabbit anti-phospho-S6 (Ser240/244) (activated S6) | D68F8 | Cell Signaling Technology, 5364 | 1:5000 |
| rabbit anti-total S6 (Ser240/244) | 5G10 | Cell Signaling Technology, 2217 | 1:1000 |
| rabbit anti-phospho-4E-BP1 (Ser65) (activated 4E-BP1) |  | Cell Signaling Technology, 9451 | 1:1000 |
| rabbit anti-total 4E-BP1 | 53H11 | Cell Signaling Technology, 9644 | 1:1000 |
| mouse anti-ACTIN | 8H10D10 | Cell Signaling Technology, 3700 | 1:10000 |
| **Secondary HRP-conjugated antibodies** | | | |
| goat anti-mouse IgG |  | Southern biotech, 1010-05 | 1:2000 |
| goat anti-rat IgG |  | Southern biotech, 3010-05 | 1:5000 |
| goat anti-rabbit IgG |  | Southern biotech, 4010-05 | 1:5000 |
